# Sparse networks of conformational fluctuations communicate signals within proteins

**DOI:** 10.1101/2025.05.28.656549

**Authors:** Kaitlin Trenfield, Milo M. Lin

## Abstract

To respond to environmental cues, proteins must amplify angstrom-scale signals across nanometers in the presence of thermal fluctuations. A prevailing view is that thermal fluctuations attenuate (*1–7*) signal-bearing coherent motions (*8–11*), yet numerous experiments correlate signaling state with fluctuations themselves (*12–36*). Here, we show that residue-level fluctuations encode “geometric bits” that are communicated within a sparse 3D network of shared entropy. We demonstrate this by developing an open-source framework that discovers shared entropy networks by inferring discrete residue conformations from molecular dynamics simulations of protein structure and finding maximum-likelihood tree distributions with minimal assumptions, enabling multiscale conformational entropy calculation without exhaustive enumeration. We validate our approach against an array of experimental data modalities probing sequence and ligand-dependent functions of PDZ and estrogen receptor ligand-binding domains, accurately predicting allosteric hotspots in saturation mutagenesis with residue-scale resolution, local entropies correlated with NMR and HDX dynamics, and global entropies in agreement with calorimetry without fitting. Comparing networks of the six human steroid receptors recovers phylogenetic history, providing evidence that evolution achieves functional diversity by reprogramming entropy within a fixed protein fold. The ability to transmit signals by harnessing thermal fluctuations categorically distinguishes proteins from human-designed communication channels, for which fluctuations are noise to be minimized.

## Introduction

To orchestrate environment-aware behavior, signaling proteins must function reliably as both receptors and transmitters in the presence of thermal fluctuations. Signals include ligand binding and post-translational modification, and their transmission modulates activities such as molecular recognition and catalysis at faraway sites, a phenomenon called allostery (*37*). Allosteric proteins have the capacity for context-dependent activity, making them basic building blocks of scalable biological circuits (*38*).

How allostery emerges from intramolecular interactions is an open question. In a prevailing view, allostery is modeled by interpreting global modes of coherent motion (*1–7*) or mean displacements (*8, 9*) as functional signals, with residual motions such as local conformational fluctuations treated as noise. This picture comports with the behavior of human-designed communication channels, well-described by communications systems theory (*39*), in which thermal fluctuations only serve as noise. However, theoretical considerations (*40, 41*), bioinformatic evidence (*42–44*), and numerous experiments suggest a central role for conformational entropy in protein allostery (*19–36*), with allosterically-responsive dynamical signatures often predicting functional specificity without significant changes in average structure (*12, 13, 28–32*).

Conformational entropy has been estimated from experimental structures (*25, 32, 45*) and NMR dynamics (*20, 21*). However, experimental methods do not resolve correlated fluctuations, in which the conformation of one residue is influenced by the conformation of other residues (*46, 47*). These correlations would give rise to shared entropy (also called mutual information) which must be subtracted to obtain accurate estimates of total entropy. Computational approaches (*48–51*) have not quantitatively predicted experimental observables such as allosteric sensitivity or conformational entropy. Consequently, we lack accurate 3D maps of allosteric networks, limiting our ability to predict responses of proteins to perturbations, fine-tune the binding affinities of designed proteins, and rationally engineer protein signaling (*52–54*). More fundamentally, it has not been established whether the couplings between localized conformational fluctuations constitute functional communication channels, or if such fluctuations merely serve as noisy readouts of information transmitted via coherent global modes of displacement and vibration.

Here, we introduce ciMIST (conformational inference-maximum information spanning tree), a framework that maps conformational ensembles observed in molecular dynamics (MD) simultions to simplified thermodynamic networks of interacting residues. Leveraging the discreteness of free energy wells in dihedral angle degrees of freedom (*50, 51, 55–57*) and sparsity of intra-protein couplings (*50, 51, 56–62*), ciMIST infers whole-residue discrete conformations from multidimensional residue statistics and then constructs the maximum likelihood spanning tree (*63–65*) as a global statistical model for the joint distribution of residue conformations. Together, these ciMIST features enable calculation of conformational entropy contributions of arbitrary residue sets, overcoming a limitation of existing information-theoretic analyses of MD and facilitating experimental predictions.

We show that ciMIST provides accurate and interpretable 3D models of entropic allostery in two well-studied protein domains, PDZ3/PSD95 and the estrogen receptor *α* ligand binding domain (ER*α*-LBD). Using all-atom, explicit solvent MD simulations of multiple liganded and mutational states of each, we demonstrate that ciMIST identifies allosteric hotspots at residue-scale precision across a saturation mutagenesis library, and predicts local entropies correlated with experimentally measured protein dynamics as well as global entropy differences consistent with calorimetry to within measurement uncertainty.

We illustrate how ciMIST provides an intuitive platform to quantitatively reveal biophysical mechanisms hidden within MD trajectories, in the case of ER*α*-LBD revealing that ligand discrimination is achieved through entropy compensation between helix formation and loop disentanglement in a disorder-prone breast cancer mutation hotspot. Extending our analysis to simulations of all six human steroid receptors, we find that comparing ciMIST-inferred entropy networks between proteins recapitulates their phylogenetic relationships, whereas traditional structure and dynamics-based methods do not. This shows how evolution, by modifying sparse shared-entropy networks, could diversify protein signaling within a family without changing the protein fold.

That ciMIST networks are sufficient to explain this large set of experimental data across the proteins studied suggests that conformational fluctuations, acting like “geometric bit flips”, are a source of sparsely-organized digital communication underlying allostery in signaling proteins. Collectively, these results illustrate how proteins work with thermal fluctuations (rather than against them) to transmit signals between precise locations, demonstrating a contrast between designed and evolved communication channels.

### ciMIST: Dynamics-based intra-protein information network

At first, inferring network models of protein conformational entropy from MD might seem impossible due to the complexity of protein energy functions. However, saturation mutagenesis reveals insensitivity of binding free energies to mutations at most amino acid positions (*58*). This suggests that the complexity of the energy as a function of atomistic positions gives way to sparsity in the free energy as a function of coarse-grained features at individual amino acid positions. We reasoned that this robustness might originate from the existence of well-separated basins in energy landscapes of amino acid dihedral angles, which set thresholds that energetic perturbations must exceed to significantly alter local geometric structures. We therefore hypothesized that protein conformational entropy would exhibit a similar effective sparsity when parameterized in terms of discrete conformations of entire residues. To make this hypothesis testable, we developed a framework for inferring whole-residue conformations and sparse conformational entropy networks from MD simulation data (figure 1).

**figure 1:**
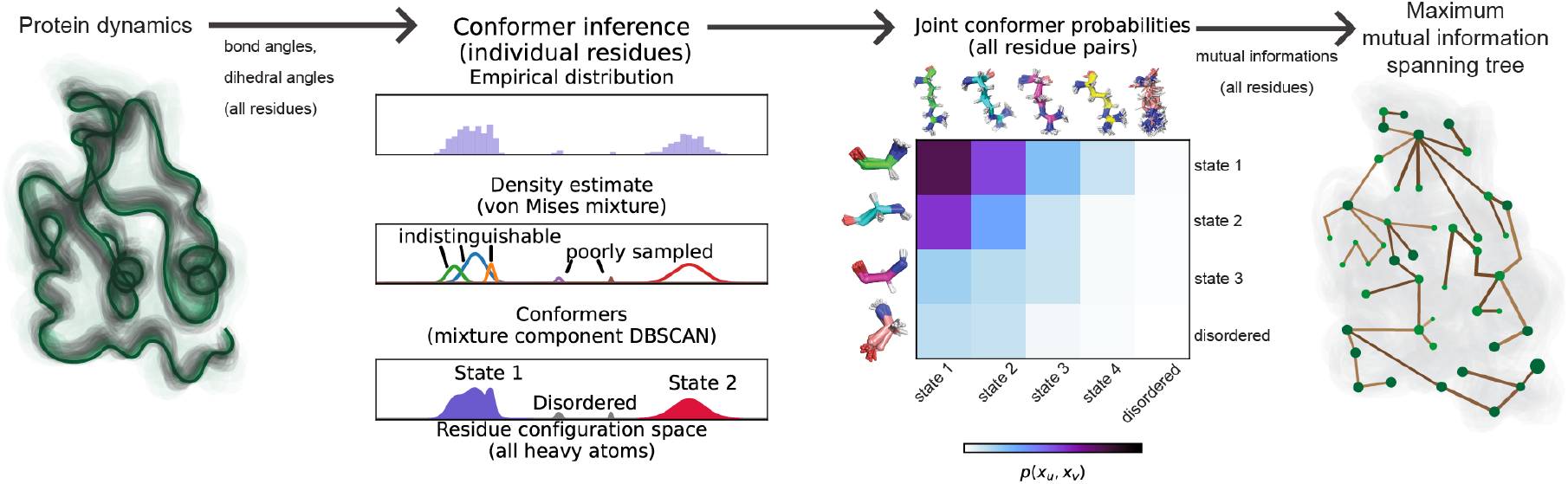
ciMIST infers maximum information spanning trees linking single-residue conformations into a global statistical model. All frames of a molecular dynamics ensemble are mapped to internal coordinates. For each residue, a separate von Mises mixture model is fit to the observed residue configurations, parameterized using bond and dihedral angles. Mixture components for each residue are coarse-grained by applying DBSCAN, which identifies residue conformations by merging components whose densities overlap, and maps all low-probability components to a single disordered state. Conformational probabilities and mutual informations are computed for each pair of residues residue, and the maximum information spanning tree is then found.

To capture network sparsity, we modeled joint distributions of residue conformations observed in MD as tree distributions. In a tree distribution, the probability of any global conformation *p* (*x*_1_…*x*_*L*_) depends only on *L* first-order probabilities { *p* (*x*_*u*_)} and *L* − 1 pairwise probabilities { *p* (*x*_*u*_, *x*_*v*_)}; the residues and pairs are interpretable as the nodes and edges (respectively) of a spanning tree. Writing *E* (*T*) for the set of pairwise interactions that directly influence global probability,

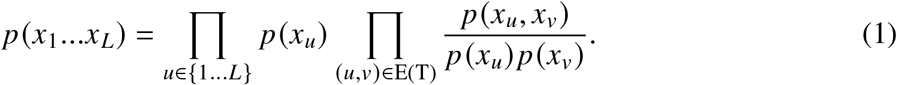

From the Gibbs-Shannon entropy formula 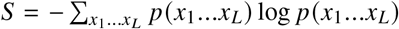, it follows that the conformational entropy is

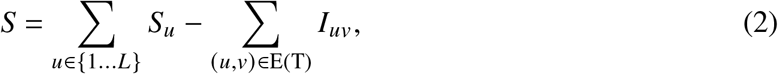

where 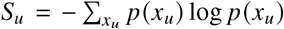 is the marginal entropy of residue *u*, E(T) denotes the set of edges in the spanning tree, and 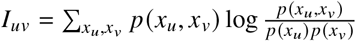 is the mutual information (MI) between residues *u* and *v*. The optimal tree approximation to any probability distribution is simply the spanning tree that maximizes 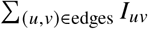, called the maximum information spanning tree (MIST) (Materials and Methods) (*65*). Determining the optimal tree structure and the conformational entropy can be done in polynomial time, requires knowing only pairwise probabilities, and does not require enumerating all possible trees or conformations. We estimate the requisite probabilities from MD simulations using empirical frequencies (Materials and Methods). Equation 2 is also decomposable into single variable contributions as

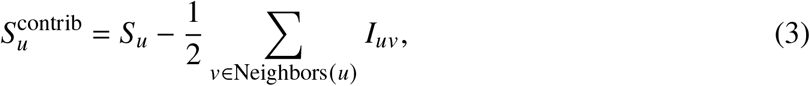

where Neighbors(*u*) denotes all *v* such that *p* (*x*_*u*_, *x*_*v*_) appears as a factor in equation 1 and the factor of 1/2 ensures MI is shared equally in accounting for contributions. Hence, by combining the MIST algorithm with equation 2, we can use a tree distribution to estimate total protein conformational entropy, while equation 3 gives a pragmatic decomposition by which to analyze local entropy.

Parameterizing conformational entropy using a tree distribution also formalizes the hypothesis that any residue pair shares information along a unique path. While this cannot be exactly true, it should be a good approximation when the information structure is sparse because a tree is the sparsest network that connects all residues. Furthermore, introducing more connections would require that we resort to enumeration or introduce numerical approximations to determine the entropy.

In devising an algorithm for inferring residue conformations, we aimed to (i) identify conformations with non-overlapping free energy wells in a multidimensional landscape and (ii) infer conformations based on statistical resolution with minimal prior assumptions. We employed a two-stage procedure (figure 1, Conformer inference; supplemental figure S2). First, we estimated continuous free energy surfaces for each residue by fitting over-parameterized von Mises mixture models to the bond and torsion angles observed in MD. Next, we used DBSCAN to cluster mixture component distributions using an overlap-based distance and a minimum cluster probability of 0.01. The use of DBSCAN to cluster overlapping components ensures conformations are distinguishable from each other and well-sampled. The same input parameters for mixture models and DBSCAN were used for all analyses in this work. Across all MD ensembles analyzed in this work, we find that residues typically have fewer than 10 conformations (supplemental figure S3). Importantly, because the number of conformations are determined by statistical resolution criteria, our approach controls the bias and variance in our entropy and MI estimates (*66*).

Using the residue-specific states found by our conformer inference algorithm, we calculated joint probabilities of occupying different conformer states for all pairs of residues, and then used the mutual information calculated for residue pairs to construct the maximum information spanning tree. The combined open source framework, which we term ciMIST (figure 1), extracts conformational entropy networks from MD ensembles. For each MD ensemble analyzed in this paper, we used the *L* − 1 pairwise probability distributions used by the optimal tree distribution to calculate the remaining pairwise distributions, verifying that the sparse set of pairwise interactions generally provides good approximations to held-out pairwise probability distributions (supplemental figure S4). To address statistical uncertainty, we initialized the program from different random seeds for each MD ensemble. We then devised an uncertainty quantification method that summarized the variability in the entropy across these runs in terms of uncertainty in the single and pairwise probability distributions and the uncertainty in the network structure (Materials and Methods, supplemental figure S7). All trees shown in this work are with entropies closest to the average of 21 runs for that ensemble, with uncertainty quantified as described in the methods section. For the remainder of this work, unless otherwise specified, we report entropy in units of contribution to the thermodynamic free energy, which is obtained by multiplying by *RT*, where *R* is the gas constant and *T* is temperature.

### A unified dynamics-based model for entropic allostery in PDZ3

We used ciMIST to model entropic allostery in the third PDZ domain from PSD95 (PDZ3) (figure 2A), a small signaling domain with a well-studied allosteric network. Mutagenesis, structure, and coevolution show how residues in the *β*2 − *β*3 loop shape specificity (*58, 59, 67*). Entropic allostery is supported by calorimetry and NMR showing how a non-conserved helix (H3) allosterically stabilizes apo-state fluctuations, reducing entropy lost when a peptide (CRIPT) binds (*28*). These insights, obtained from different experimental approaches with varying levels of quantitative resolution, provide a patchwork picture of allosteric sensitivity in PDZ. Based on simulated dynamics beginning with crystal structures, we first used ciMIST to reproduce known properties into a unified quantitative model. We then challenged ciMIST to provide new insights about allosteric coupling within PDZ: inferring residue-scale responses to perturbations as well as generating whole-protein changes in conformational entropy from the sum of the parts and their interactions.

**figure 2:**
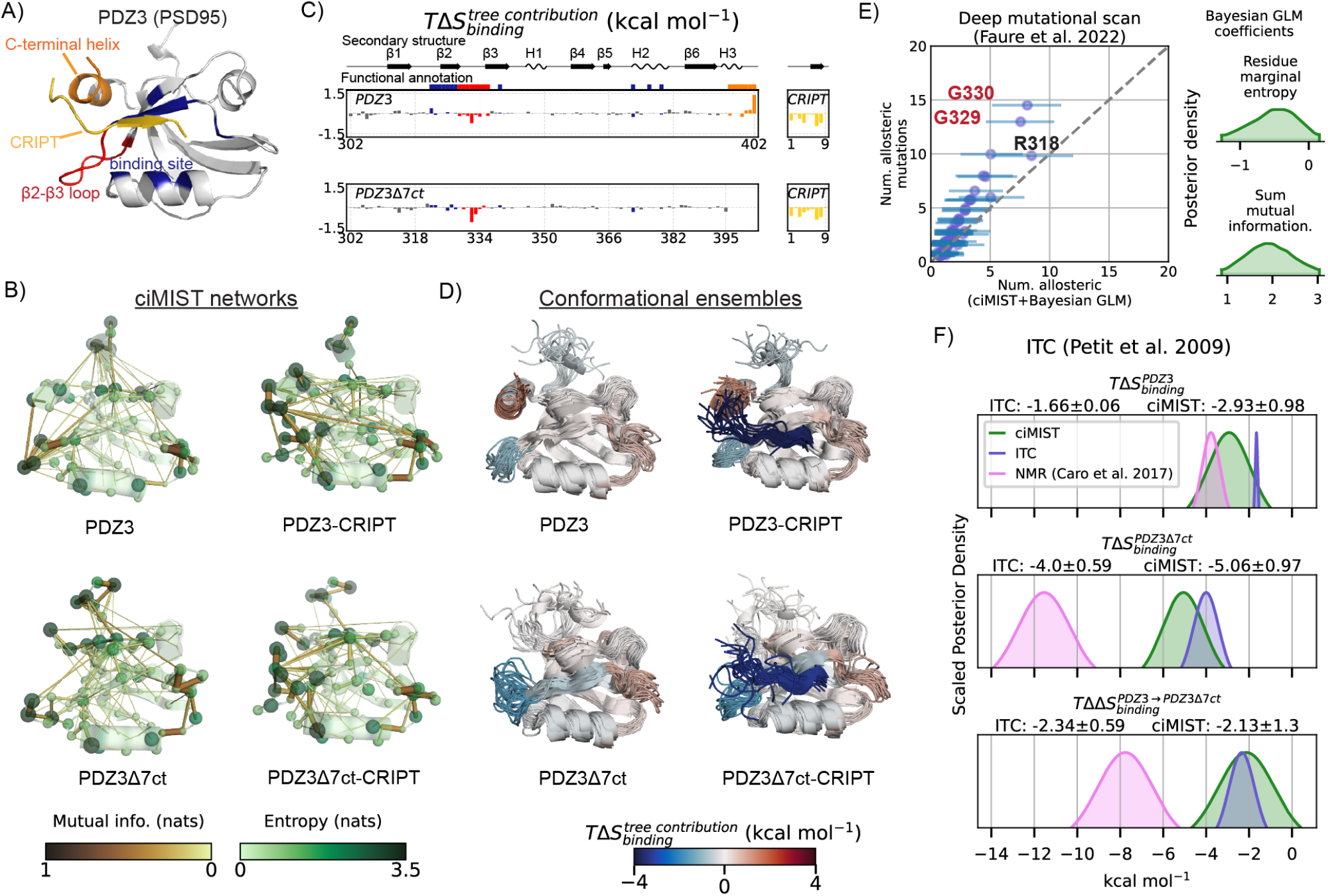
Connecting molecular dynamics to experimental signatures of allostery in PDZ3. A) Structure of the third PDZ domain (PDZ3) from PSD95 bound to the CRIPT peptide. B) Maximum information spanning trees for PDZ3 (top) and PDZ3Δ7ct in apo (left) and CRIPT-bound (right) states. Sphere volumes of C*α* atoms (shades of green) are proportional to residue marginal entropies, while volumes of links (shades of brown) are proportional to mutual information. C) Contributions of individual residues to the conformational entropy difference between apo and CRIPT-bound states. Regions of of the protein are color-coded as in (A). D) 20 randomly-selected snapshots from simulations of PDZ3 (top) and PDZ3Δ7ct (bottom) with secondary structural elements colored by contribution to the binding conformational entropy. CRIPT is classified as a single element. E) Prediction of allosteric mutational propensities in saturation mutagenesis from ciMIST residue thermodynamic features using a Bayesian GLM. Residue marginal entropies and summed mutual informations with tree neighbors from the CRIPT-bound state of PDZ3Δ7-ct were used as predictors. F)Posterior densities for whole-protein conformational entropy differences calculated by ciMIST, entropies measured with ITC (*28*), and conformational entropy differences estimated by NMR (*21*). ITC and NMR posteriors are Gaussians with reported means and standard deviations. *T* = 298*K*.

We performed MD simulations (*68, 69*) (Materials and Methods, Supplemental Table S1) of PDZ3^302−402^ and (henceforth: PDZ3) and PDZ3^302−395^ (PDZ3Δ7ct, lacking H3) in apo and CRIPT-bound states. We also simulated the isolated CRIPT peptide. This constitutes exhaustive simulation of the experimentally characterized thermodynamic cycle that revealed H3-mediated entropic control of the PDZ3-CRIPT interaction (*28*).

The ciMIST networks of these simulations show that the *β*1 − *β*2 and *β*2 − *β*3 loops act as information network hubs (figure 2B), highlighting known allosteric sites. Central to the *β*1−*β*2 loop hub is ARG318, which is known to modulate binding kinetics in response to ionic strength (*70*). In the *β*2 − *β*3 loop, GLY330 shows high MI with many residues, consistent with its known mutational sensitivity (*58,59,67*) and the role of this site in determining specificity between peptide classes (*67*).

Using equation 3, we quantified contributions of regions of PDZ3 to conformational entropy changes (figure 2 C-D). As expected, CRIPT stabilization imposes a baseline entropic binding penalty: −3.35 kcal mol^−1^ upon binding to PDZ3Δ7−ct and −3.56 kcal mol^−1^ upon binding to PDZ3. PDZ3 partially offsets this with an increase of 1.63 kcal mol^−1^ in entropy contributed by C-terminal residues 399-402 (figure 2 C). The *β*2 − *β*3 loop alo contributes a loss of entropy which is greater for PDZ3Δ7ct (−1.27 kcal mol^−1^) than PDZ3 (−0.76 kcal mol^−1^) due to loss of pre-stabilization by H3.

To quantitatively test whether mutual information predicts allosteric hotspots, we used ciMIST residue-level thermodynamic features to build a Bayesian generalized linear model (GLM) (Materials and Methods) of allosteric mutational sensitivities in a deep mutational scan (*58*). Using the marginal entropy and the sum of MI with network neighbors from the tree for PDZ3Δ7-ct, we obtained excellent correlation with reported site propensities for allosteric mutations (figure 2E). Posterior distributions showed that neighbor-summed MI positively correlated with allosteric mutational propensity (95% credible interval: [0.70, 2.61]). The posterior distribution for the coefficient of residue marginal entropy concentrated on negative values (95% credible interval: [−1.30, 0.20]), indicating that cooperative residue dynamics, rather than mere flexibilities, predict allosteric function.

Conformational entropy differences predicted by ciMIST agree with measured binding entropies of PDZ3 and PDZ3Δ7-ct (figure 2F). Differences in residue entropy contributions also correlate significantly with reported changes in NMR backbone order parameters between the apo states (Pearson *r* = −0.69, *p* < 10^−8^), but not side-chain order parameters (*r* = −0.09, *p* = 0.67), (supplemental figure S5). This partially supports the model, suggested based on NMR, which attributes the change in binding entropies to differences in apo-state ensembles, with the main difference being that our analysis supports no special role for side chain entropies. Notably, our calculations are in better quantitative agreement with calorimetric entropies than the NMR entropy meter, which estimates conformational entropy using only side chain order parameters (*21*).

### A unified model for entropic allostery in estrogen receptor *α*

We next used ciMIST to model entropic allostery in ER*α*-LBD, a larger signaing protein domain for which allosteric regulation shapes outcomes in multiple cell types, including as the primary determinant of breast cancer cell fate (*71, 72*). Published data suggested a number of possible roles conformational entropy might play in its allosteric regulation. First, a loop region spanning helices 9 and 10, dubbed the H9-H10 instability, is unresolved in most experimental structures but is a hotspot for breast cancer mutations. Second, despite over 300 ER*α*-LBD structures in the PDB, the structure of its apo state is unknown, leaving a fundamental gap in understanding of how the “off” state is maintained in healthy conditions and disrupted by breast cancer mutations such as Y537S. What is known is that binding of ligands leads to globally decreased hydrogen-deuterium exchange (HDX) (*12*) and entropy losses exceeding 10 kcal mol^−1^ (*73*).

We applied ciMIST to simulations of ER*α*-LBD to elucidate the relationship between these allosterically-sensitive indicators of disorder and conformational entropy. For all liganded and mutational states here, we simulated ER*α*-LBD homodimers, reflecting nanomolar-scale homodimer affinity (*74*) and micromolar-scale receptor concentrations in experimental datasets (*12, 13, 73*). For each thermodynamic state, we again simulated six replicates and concatenated these to obtain a single conformational ensemble (Materials and Methods, supplemental table S1). We concatenated the ensembles of individual subunits, so that all conformational distributions and couplings are averaged over the two copies in the homodimer, improving statistical resolution at the cost of explicitly capturing inter-domain cooperativity.

### Predicting the effects of thermostabilizing mutations on ERα binding entropies

We first used ciMIST to compare human ER*α*-LBD to an engineered thermostable estrogen receptor (prsER*α*, figure 3A) (*73*). prsER*α* was designed by introducing 22 PROSS-suggested (*75*) substitutions into the ER*α*-LBD sequence outside of major allosteric sites. While conserving binding affinities and ligand-dependent structure, prsER*α* displays altered binding entropy for genistein (GEN), a phytosterol present in human diet that is allosterically discriminated from estradiol by human ER*α* (*74*). Hypothesizing that altered binding entropy was due to stabilizing mutations in H10, we simulated both receptors bound to estradiol and genistein.

**figure 3:**
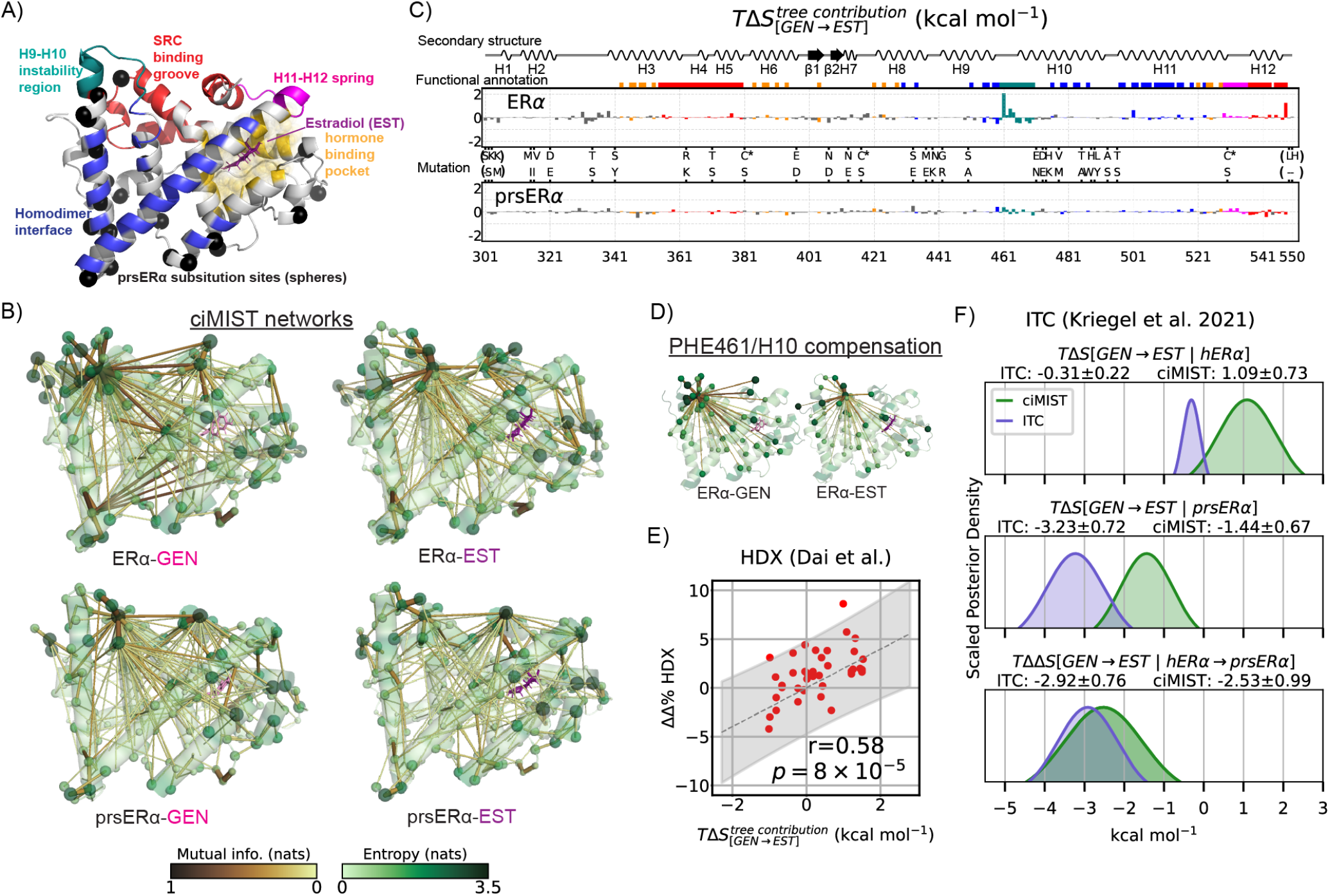
Entropic effects of 22 stabilizing mutations in ER*α*. A) View of a single subunit of ER*α* with functional annotation. B) ciMIST networks for the ER*α* (top) and a prsER*α* (bottom) bound to genistein (left, pink sticks) and estradiol (right, purple sticks). Sphere volumes of C*α* atoms (shades of green) are proportional to residue marginal entropies, while link volumes (shades of brown) are proportional to mutual information. Note that while simulations were of homodimers, ciMIST was applied to the aggregate statistics of both subunits. C) Differences in residue entropy contributions between genistein- and estradiol-bound states for ER*α* (top) and the prsER*α* (bottom). C^∗^ denotes positions of cysteine-serine substitutions made for calorimetry (*73*) but not in our simulations. Parentheses denote positions not present in the calorimetry experiment. Regions are color-coded as in (A) D) Connections in ciMIST networks between PHE461 and other residues for human ER*α* ensembles. E) Correlation between differences in contributions of protein segments to total conformational entropy and average percent differences in deuteration levels (*12, 13*) between estradiol and genistein-bound states of human ER*α*. F) Estimated posterior densities of entropy differences from ITC (*73*) and calculated using ciMIST. Uncertainties for ITC are estimated using the standard deviation of three replicates provided by the authors. *T* = 298*K*.

The H9-H10 loop region is an information hub in the ciMIST networks for all four ensembles, with PHE461 especially prominent (figure 3B-D). In both human receptor trees, PHE461 displays high mutual information with H9-H10 loop residues, H10 residues, and with TYR459 in H9, but with greater MI when ER*α*-LBD is bound to genistein (relative to estradiol). Driven by increased MI, PHE461 contributes 2.06 kcal mol^−1^ less entropy in the genistein-bound state, even though the contribution from marginal entropy differs only by 0.1 kcal mol^−1^ . Interestingly, H10 has less helical order in the genistein-bound state (supplemental figure S8). Thus, the cost of helix formation in the estradiol-bound state is compensated by increased H9-H10 loop entropy due to disentanglement of PHE461 from H10. Supporting our hypothesis, we found similar couplings in prsER*α* trees, but mutational pre-stabilization decreased marginal entropies and MI in this area, greatly attenuating differences in PHE461/H10 entropic compensation.

To test the agreement between local entropies and experimental dynamics, we compared differences in the entropy contributions of segments (computed by summing equation 3 over segments) with differences in hydrogen-deuterium exchange (HDX) between estradiol and genistein-bound states of ER*α*-LBD (*13*). We found significant correlation (Pearson *r* = 0.58, *p* = 8 ×10^−5^) between these (figure 3E), validating that changes in local entropies capture changes found in solution-state experimental dynamics. Entropy contribution differences correlated better with HDX changes than summed residue marginal entropy differences (Pearson *r* = 0.31, *p* = 0.052), supporting that the experimental dynamics are intrinsically cooperative.

We also found that ciMIST predicted entropy differences between estradiol and genistein-bound MD ensembles are within 1-2 kcal mol^−1^ of experimental values (*73*) for both ER*α*-LBD and prsER*α*, although exact agreement should not be expected here because our method estimates only protein conformational entropies without accounting for differences in ligand, solvent, or vibrational entropies within conformational states (figure 3F). However, our calculated difference in how the two proteins respond to estradiol compared to genistein (*T*ΔΔ*S*) falls within statistical uncertainty of experiment (figure 3F), as expected if differences attributable to non-conformational sources are similar for the two receptors.

By using ciMIST to compare ER*α*-LBD to prsER*α*, we found a ligand-dependent entropic compensation mechanism in the H9-H10 instability region of ER*α*-LBD allosterically discriminates between similar ligands. Quantitative agreement between ciMIST predictions and two independent experimental datasets strengthens confidence in this mechanism.

### Rewiring of the ERα allosteric network in the Y537S mutant

We next wanted to know whether ciMIST could identify significant differences between apo-state conformational ensembles of wild-type ER*α*-LBD and the constituitively-active Y537S somatic breast cancer mutant. The Y537S mutant drives estrogen-independent proliferation of breast cancer cells and resistance to anti-estrogen therapies (*15, 71, 76*). We applied ciMIST to simulations of apo states of ER*α*_*Y*537*S*_ and wild-type ER*α*. For the wild-type, we used homology modeling to create a starting structure for MD simulations using structures of the Y537S and D538G mutants in their apo states as templates (Materials and Methods). H12 remained oriented towards the dimer interface (agonist conformation) in both ensemble-averaged structures (figure 4A).

**figure 4:**
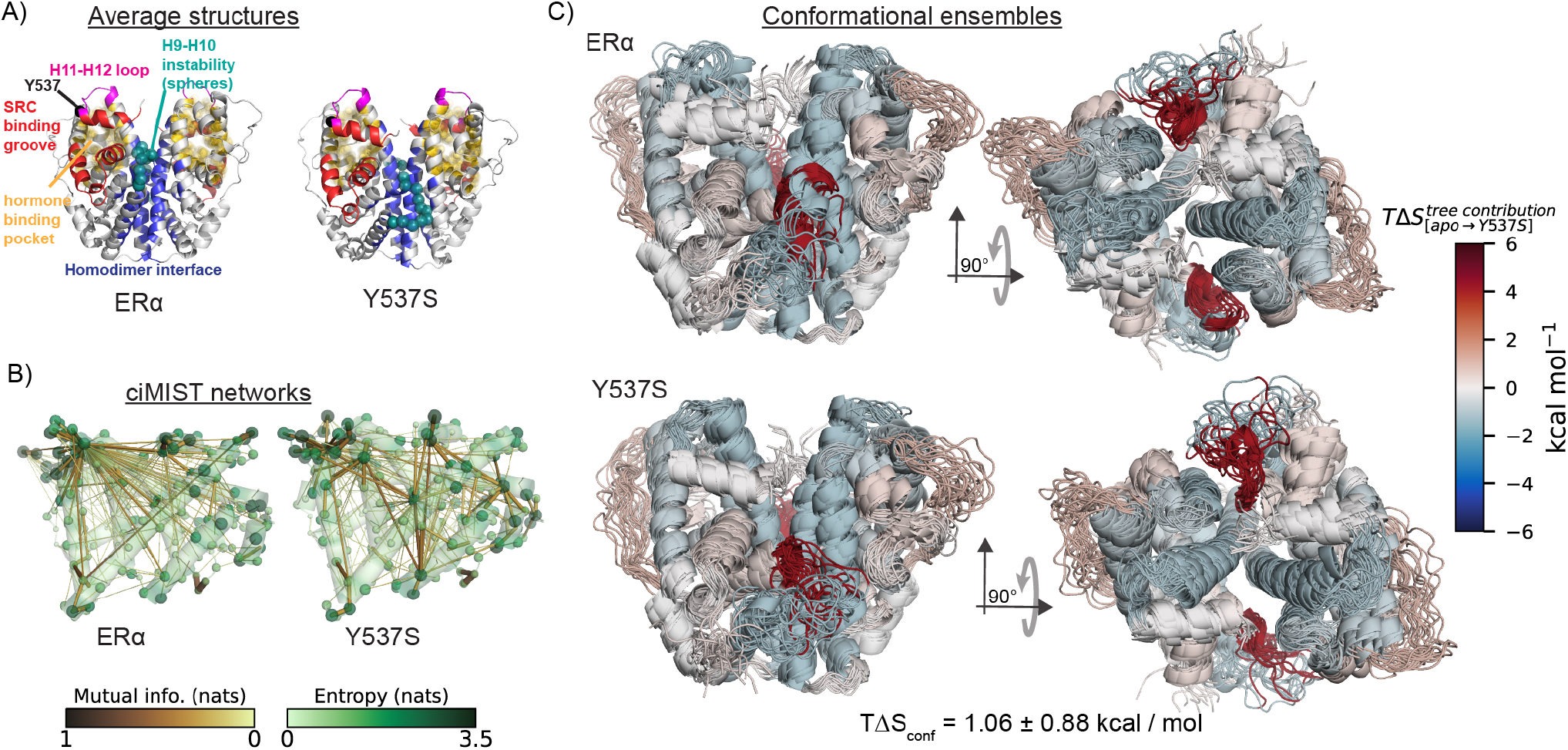
Redistribution of entropy in the unliganded (apo) state of the Y537S estrogen receptor *α*. A) Average structure observed in simulations of human ER*α*-LBD homodimer and the Y537s variant. Structures were computed by superimposing all frames onto the first and then computing the average positions of atoms over all frames. B) Maximum information spanning trees for the unliganded wild-type and Y537S variants of ER*α*. C) 20 randomly selected snapshots from MD simulations of unliganded wild-type and Y537S variants of ER*α*. Secondary structural elements are colored by their contributions to the difference in entropy between mutant and wild-type states, *T* (*S*_Y537S_ − *S*_wild-type_). *T*Δ*S*_*conf*_ is the ciMIST calculation of difference in conformational entropy of Y537S from the wild-type. *T* = 298*K*.

The wild-type ciMIST network features mutual information between PHE461 and binding site residues HIS524 and GLU353 (figure 4B, supplemental figure S6D), indicating intrinsic allosteric sensitivity of PHE461 to binding site conformations. The ciMIST network of the Y537S mutant was noticeably different, consistent with altered allostery (figure 4B); in particular, many PHE461 connections were weakened or lost, while LYS472 and ARG434 emerged as new information hubs.

Relative to wild-type, Y537S displayed increased PHE461 entropy contribution (from decreased PHE461-H10 coupling) and decreased H10 entropy contribution (from increased LYS472-H10 coupling). We also observed decreased entropy contributed by H3, H11, binding pocket residues, and the H11-H12 loop (figure 4C, supplemental figure S9C), consistent with *in vivo* Y537S activities comparable to those of the estradiol-bound wild-type (*15*). Conformational stabilization of the binding site and H11-H12 in Y537S correlated with decreased hormone-binding pocket hydration (Supplementary figure S6A-B). ciMIST predicts similar conformational entropies for these ensembles, suggesting that cooperative stabilization of H11-H12 and the protein core (figure 4C), rather than global stabilization, might confer hormone-independent crystallization and signaling activities.

### Local entropy compensation and global disorder in ERα regulation

Aiming to build a comprehensive model ER*α*-LBD accounting for local and global conformational instabilities, we applied ciMIST to simulations of ER*α* bound to 4-hydroxy-tamoxifen and lasofoxifene (supplemental table S1), both of which inhibit estrogen-dependent ER*α* activities by reorienting H12 away from the homodimer interface (antagonist H12 conformation) (*71*). Combined with aforementioned simulations of ER*α*-LBD, this provided a balanced sampling of all known ER*α*-LBD folds. Protein conformational entropy differences from the apo state are between −8 and −14 kcal mol^−1^ for all liganded states (figure 5A) – in the vicinity of the experimental binding entropies (*73*).

**figure 5:**
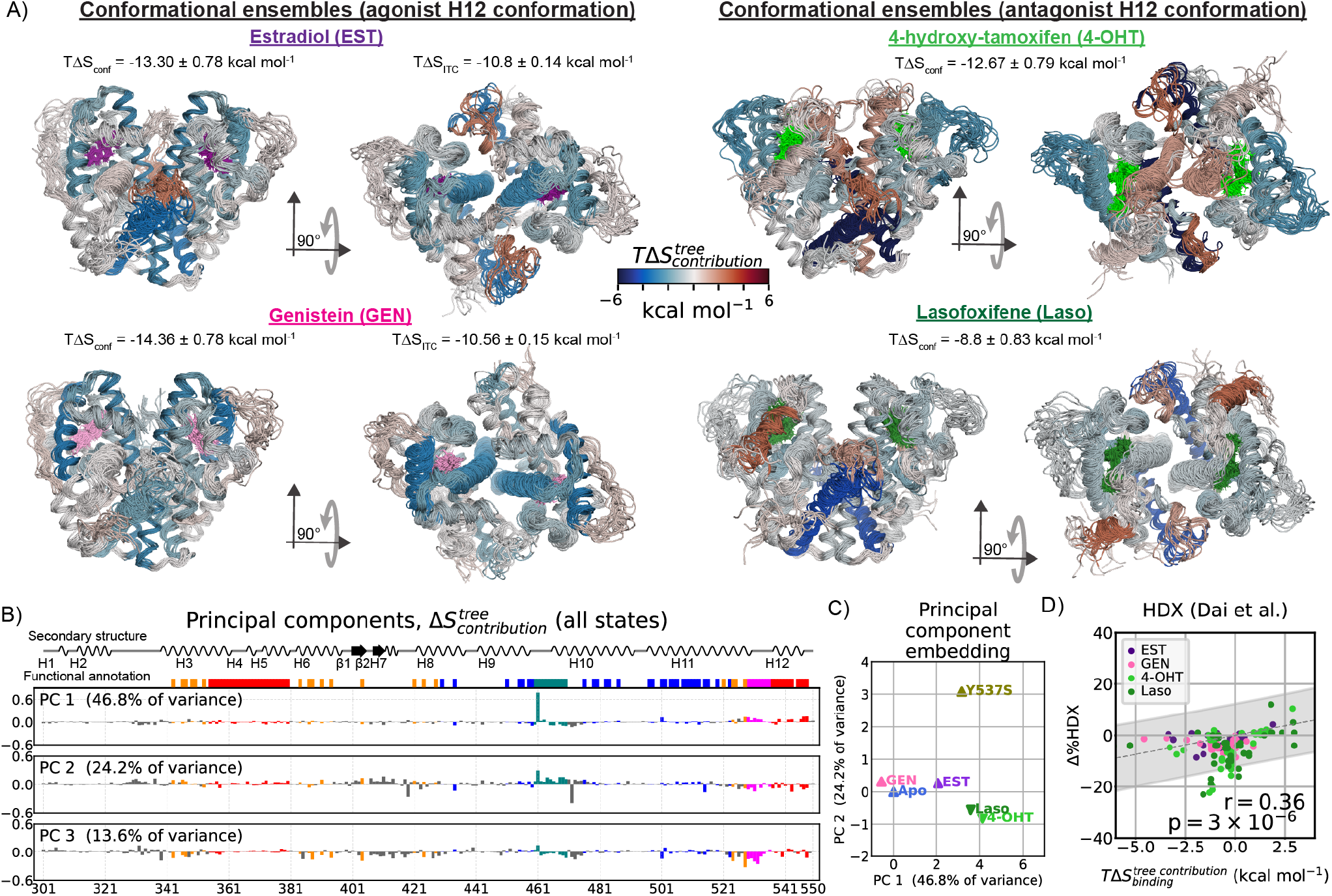
Entropic allostery in estrogen receptor *α*. A) 20 randomly selected snapshots from simulations of ER*α*-LBD homodimers bound to estradiol (purple), 4-hydroxytamoxifen (lime green), genistein (pink), lasofoxifene (forest green). Secondary structural elements are colored by their contribution to the difference in entropy from the wild-type apo state. *T*Δ*S*_*con f*_ is the ciMIST calculation of change in conformational entropy relative to the apo state, with posterior standard deviation as the stated uncertainty. *T*Δ*S*_*ITC*_ denotes experimentally-measured binding entropy (*73*), for which uncertainties are standard deviations of three replicates. B) First three principal components of changes in entropy contributions of the set of liganded states shown here as well as the Y537S apo state from the wild-type apo state. C) Projection of residue entropy contribution vectors for ER*α* states onto the first two principal components. D) Correlation between average contributions of residues in protein segments to binding conformational entropy and average percent difference in deuteration levels between apo and liganded states (*12, 13*). The gray band demarcates 95% prediction intervals. *T* = 298*K*.

We applied principal component analysis to differences in residue entropy contributions between all ER*α* ensembles and the wild-type apo state (figure 5B-C). The leading principal component, accounting for 46.8 percent of variance in entropy contributions, captures PHE461/H10 entropy compensation, validating this as a general mechanism of ER*α* allosteric regulation (figure 5B) modulated by agonist and antagonist H12 conformations. The second component separates antagonist ligands from apo-Y537S, consistent with resistance of Y537S to anti-estrogen therapies (*15, 71*). The third component captures stabilization of the protein core (figure 5B, supplemental figure S6C).

We compared entropy contribution differences to changes in HDX (*12, 13*), this time using the wild-type apo state as a baseline. We found significant correlation (Pearson *r* = 0.36, *p* = 3 × 10^−6^) between entropy contribution and HDX differences (figure 5D) and weaker correlation using summed residue marginal entropies (Pearson *r* = 0.3, *p* = 0.0001). While HDX is interpreted as measuring dynamics in ER*α* (*12, 13, 15, 77*), to our knowledge this is the first study to bridge the gap between HDX and calorimetry using an MD ensemble. Notably, neither our sampled MD ensembles nor ciMIST parameters were fit to experimental data.

These models quantitatively explain multiple observations – large losses of entropy (*73*) and hydrogen-deuterium exchange (*12, 13*) upon ligand binding – as consequences of global conformational entropy loss, while also revealing ligand discrimination by PHE461/H10 entropy compensation. Further supporting entropic compensation are increased homodimerization and estrogen-independent activities conferred by somatic breast cancer mutations F461V and S463P (*78*). The H9-H10 region is also a site of proteolytic cleavage (*79*), suggesting ligand-dependent ER*α* PHE461/H10 entropic compensation may regulate ER*α* abundance. Thus, our approach may be valuable towards the design of therapeutics that accelerate receptor degradation through targeted conformational destabilization while maintaining strong antagonist positioning (*77*). Despite the size and complexity of this signaling domain, the ciMIST approach provided accurate models of entropic allostery while also uncovering a novel thermodynamic mechanism.

### Decoding the phylogenetic history of the steroid receptor family from residue-resolution entropy maps

The steroid receptor family consists of estrogen receptors *α* and *β*, and four ketosteroid receptors: progesterone, androgen, glucocorticoid and mineralocorticoid receptors. Despite adopting similar structures in the presence of their activating hormones, their allosteric mechanisms have diverged. While all active transcriptional complexes are dimers, the interface is not conserved, and thermodynamic cooperativity is observed at different stages of transcriptional complex assembly (*80–82*). Past MD analysis of dynamic contact paths to helix 12 support a mixture of conservation and divergence in ketosteroid receptor dynamics (*83*), but an analysis without prior assumption of an allosteric site and which includes estrogen receptors has not been done.

Having characterized the ER*α*-LBD signaling network, we ran MD simulations of the ligand binding domains of the five remaining steroid receptors, each bound to an endogenous agonist hormone, and inferred trees with ciMIST (figure 6A). ER*β* was simulated as a homodimer, while all ketosteroid receptors were simulated as monomers, consistent with solution-state measurements showing dimerization affinities greatly exceeding physiological receptor concentrations for hormone-bound ketosteroid receptors, but not estrogen receptors (*80–82*).

**figure 6:**
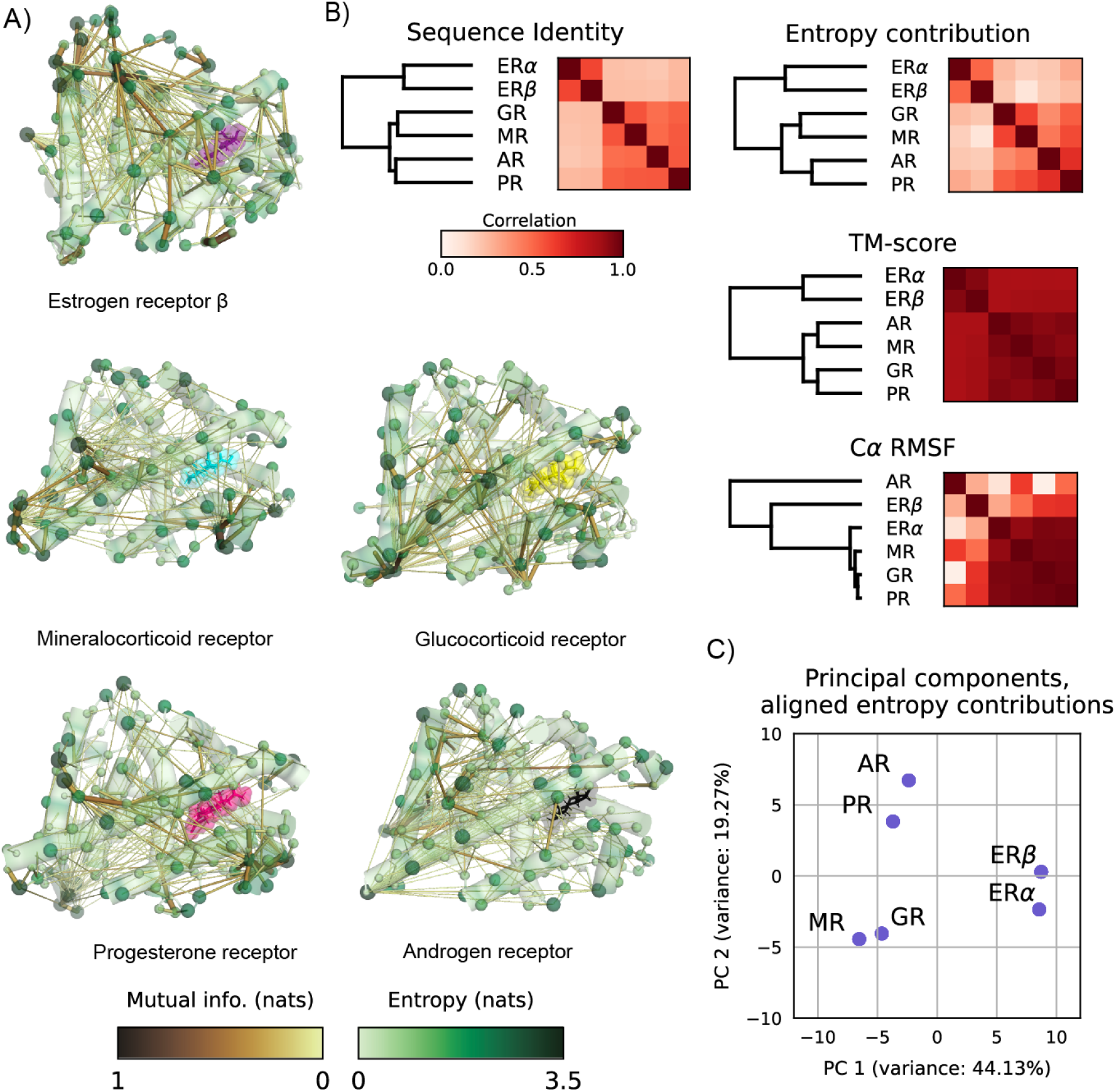
ciMIST entropy networks capture evolutionary divergence between human steroid receptor ligand-binding domains. A) Maximum information spanning trees for human steroid receptor ligand-binding domains bound to endogenous ligands. Structures are oriented as ER*α* is in figure 3A. Sphere volumes of C*α* atoms (shades of green) are proportional to residue marginal entropies, while volumes of links (shades of brown) are proportional to mutual information. B) Human steroid receptor phylogeny and application of the UPGMA clustering algorithm to similarity matrices based on sequence, ciMIST entropy contribution, structural alignment (TM-score), and dynamics (C*α* RMSF) calculated across human steroid receptors. C) The first two principal components of the aligned entropy contributions.

We used biotite (*84*) to align sequences and build a phylogenetic tree using the UPGMA algorithm, recovering the accepted phylogeny as expected (*85, 86*). Then, using the sequence alignment to align the vectors of ciMIST-computed residue entropy contributions, we calculated Pearson correlations between residue entropy contributions for all receptor-pairs and applied the same UPGMA algorithm used on sequence data to infer relationships between the receptors based on differences in their entropy maps. Residue entropy contributions diverged in a manner that mirrored evolutionary divergences. In contrast, the phylogenetic tree was not captured by root-mean-squared-fluctuations (RMSF) nor by template modeling scores (TM-scores) of the crystal structures (figure 6B).

We also found that principal component analysis of aligned entropy contributions separated receptors by phylogeny along the first two components (figure 6C). The first principal component (44% of variance) separates estrogen from ketosteroid receptors, while the second component (19% of variance) separates corticosteroid receptors from androgen and progesterone receptors – capturing both evolutionary transitions (*85*).

These results hint at a general principle: divergent information-processing networks reflect the evolutionary history of novel functional adaptations even as protein tertiary structure is conserved. Such divergence may be as necessary to precise steroidal coordination of development as are steroid-binding specificities themselves: just as estrogen signaling must differentiate estrogens from related ligands, active receptors must be distinguishable from each other to enact distinct transcriptional programs. Consistent with this, we find principal components of the entropy contributions reflect functional reprogramming at evolutionary branch points.

## Conclusions and Implications

We have shown that sparse networks of residue-level conformational fluctuations observed in microseconds of MD simulations can provide unified models of entropy-mediated allostery. While ciMIST networks predicted experimental observables with quantitative accuracy, it was equally important that they made it possible to reason about relationships between MD ensembles and protein thermodynamics in an intuitive manner. This is exemplified in the present work by our discovery of the entropic compensation mechanism in the ER*α*-LBD. By making quantitative predictions that are interpretable in terms of biochemical mechanisms, ciMIST creates the possibility of building thermodynamic models from MD ensembles that can both test pre-existing hypotheses and suggest new ones. Along these lines, we have found ciMIST to be a valuable aid towards interpreting MD ensembles and trajectories – in effect, providing a powerful lens through which to view the information structure hidden in MD ensembles.

Residues seem to communicate their conformations through networks that are sparse enough to be well-approximated by a model in which residue pairs communicate along unique paths. The discreteness of residue conformations and the sparsity of the network are, we think, closely connected. Effectively discrete states emerge from free energy wells in the multidimensional landscapes of residues’ bond and dihedral angles. Flipping these geometric bits therefore requires energetic perturbations in excess of minimum thresholds. Similar to how binding function is insensitive to mutational perturbations at most sites, conformations at most sites are insensitive to bit flips at most other sites, giving rise to sparsity.

Because mutual information contributes negatively to the entropy, residues with high mutual information can be conformationally stabilized at lower thermodynamic cost than independently fluctuating ones. The primary mutual information hubs in both PDZ3 and ER*α*-LBD were not at binding interfaces, but proximal to them. This hints at a general mechanism of entropic allostery whereby frustrated interactions in loop regions are relieved upon binding, promoting thermodynamic stability by decreasing mutual information and transmitting signal to distal sites through the engagement of pre-existing thermodynamic couplings. This mechanism requires proteins to have intrinsic sources of thermodynamic instability, as seen in our comparison of evolved and engineered estrogen receptors. Modern protein design tools tend to produce proteins with high thermodynamic and structural stability (*87*). Targeted re-introduction of thermodynamic instabilities into engineered proteins might therefore drive down binding affinities while increasing allosteric potential.

The predictive power of our features across multiple datasets and proteins indicates that the cooperative fluctuations in residue conformations we resolve are not noise, but encodings of function in the MD ensembles. One can ask more generally: is there a limit to how finely proteins can distinguish conformational ensembles? Allosteric proteins appear to have evolved through heightened selection pressures (*88*). Differential activities of transcription factors are often not explained by DNA-binding thermodynamics alone, suggesting that reliable cell fate decisions may depend on subtle differences in free energy landscapes, such as those that control kinetics (*16, 17, 89*). Proteins may therefore utilize highly sophisticated communication protocols, which information-theoretic considerations suggest should appear cryptic to naive observers (*90*). We anticipate that approaches such as ours, which connect statistical regularities of equilibrium fluctuations to functional predictions, will further decrypt the conformational codes of proteins.

## Acknowledgments

The authors would like to thank Madhav Mani, Kimberly Reynolds, Rama Ranganathan, Prashant Mishra, Seppe Kuehn, Roland Dunbrack, and Marielle Russo for feedback that significantly improved the manuscript, and Mark Kriegel and Yves Muller for sharing raw data on individual calorimetry replicates.

## Author contributions

K.T. and M.M.L. conceived the work, K.T. performed simulations and analysis, K.T. and M.M.L wrote the paper.

## Data and materials availability

ciMIST available at https://github.com/justktln2/ciMIST.

## Supplementary materials

Materials and Methods

Supplementary Text

Figs. S1 to S9

Table S1

References *(7-106)*

## Supplementary Materials for

### Materials and Methods

#### Molecular dynamics simulations

All simulations were performed using OpenMM7 using the Amberff14SB force field (*68*), TIP3P water model (*91*), Langevin integrator (*92*) with a 2 fs timestep, and a Monte Carlo barostat using Tesla V100 32GB GPUs on the UT Southwestern high performance computing cluster. Ligands were parameterized using the GAFF in Antechamber (*93, 94*). All systems were solvated in boxes with a minimum of 1.5 nanometer separation between solute and box edge. Neutralizing ions were added, followed by additional sodium and potassium ions chosen to approximate a 50 mM NaCl buffer for all PDZ domain and CRIPT simulations and a 170 mM NaCl buffer for all steroid receptors. All systems were energy-minimized using OpenMM default settings, subject to a harmonic restraint of 100 kcal mol^−1^ on positions of all alpha carbons. Equilibration proceeded by heating at constant volume from 98K to 298K while relaxing harmonic restraints to zero at a constant rate over the course of 1 nanosecond, followed by an equilibration time of 100 nanoseconds in an NPT ensemble. All PDZ domain and CRIPT simulations were sampled at 1 nanosecond intervals. Steroid receptor simulations were sampled at 0.5 nanosecond intervals. Parallel trajectories were run for each system and concatenated into a single trajectory prior to analysis. Details of simulations, including simulation times, are provided in Supplemental Table S1.

#### Inferring residue states by clustering von Mises mixture model components with DBSCAN

We define a configuration of a residue as a vector, θ = [*θ*_1_…*θ*_*M*_]^*T*^, whose entries are the bond angles and torsion angles (expressed in units of radians) needed to specify the positions of atoms belonging to that residue. We used both bond and torsion angles in our analysis because we found that setting bond angles equal to their average values distorted global protein structures upon converting from internal coordinates (lengths, bond angles, torsion angles) to Cartesian. On the other hand, setting bond lengths equal to their average resulted in very little distortion; therefore, we did not use these to define residue configurations.

For each residue, our data are an ensemble {θ_1_…θ_*N*_} of *N* residue configurations obtained by concatenating frames from all MD replicates for that thermodynamic state (combination of protein and ligand). We model the probability density as a function of residue configuration θ using a mixture of von Mises distributions:

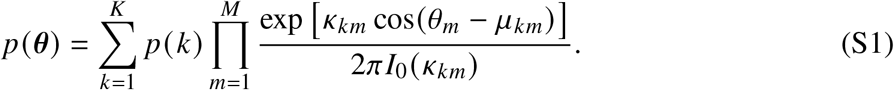

Here, the sum is over the set of *K* mixture components, *p* (*k*) is the posterior probability of the *k*-th component (with 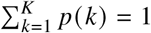), the product is over the *M* bond and torsion angles used to place the residue’s atoms, *θ*_*m*_ represents the *m*-th angle (from the aforementioned set), *µ*_*km*_ and *κ*_*km*_ denote the mode and precision parameters (respectively) of the *k*-th von Mises component along the *m*-th angular coordinate, and 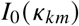 is a modified Bessel function of order 0 guaranteeing independent normalization of each component along each dimension. Note that the value of *M* depends on the residue type.

We note that each of the *K* component distributions is a product of one-dimensional von Mises distributions. Although this means that fluctuations about the individual component modes are treated as independent, the use of multiple mixture components allows this model to capture irregularly-shaped probability densities.

We chose *K* = 100 (independently of residue type) with the aim of choosing a number of mixture components greatly exceeding the number of modes of each distribution, so that the mixture models operate in the regime of density estimation, rather than peak detection.

To initialize parameters, we seeded von Mises modes using the *k*-means^++^ algorithm (*95*) with a cosine distance between individual configurations. We initialize precision parameters using random samples from uniform distributions.

Fitting proceeded by iterating the expectation-maximization algorithm until the log-likelihood function converged to within 0.01 nats per observation, obtaining distinct mixture models of the configurational probability density for each residue in the protein.

To arrive at residue states adapted to the observed probability density, rather by a predetermined selection of the number of clusters, we applied DBSCAN to the mixture components. We define the distance between components *i* and *j* in terms of the overlap between their probability densities:

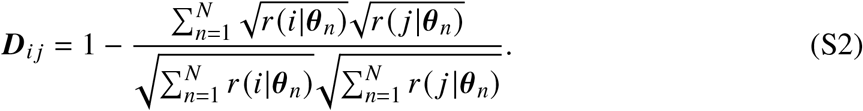

Here, *r* (*k*|θ_*n*_) is the posterior probability that observation θ_*n*_ was generated by the *k*-th mixture component (sometimes called the “responsibility” of component *k* for observation *n*), assuming the mixture model is the generative model for the data. These posterior probabilities are calculated from the component likelihoods using Bayes’ theorem. The sum runs over all *N* frames in the MD ensemble. This distance function, which is related to the Bhattacharya divergence and the Hellinger distance, takes on a value of 0 when *k* and *l* index the same mixture component and approaches a limiting value of 1 as two distributions become more sharply concentrated on distinct points. The quantity 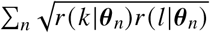, which measures the overlap between distributions, can be informally interpreted as estimating the number of observations of “mixed states” in the MD ensemble – snapshots which could have been generated by sampling from either component. Thus, clustering components using this distance metric reduces the number of observations which cannot be unambiguously assigned to a single state.

For each residue, we set the distance parameter for DBSCAN to one minus the standard deviation of all pairwise overlaps, 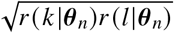. We found that this resulted in sensible partitions of the configurational spaces by inspecting lower-dimensional marginal distributions.

We set the minimum mass parameter in DBSCAN to 0.01. In the present application, this corresponds to a minimum probability mass required to classify a region of continuously high probability density as a conformation. The number of observations for all of our trajectories was greater than 10, 000, so this requirement guarantees that for any pair of residues, the expected number of counts, ⟨*n* (*x*_*u*_, *x*_*v*_) ⟩, is greater than 1 for any pair of conformations (*x*_*u*_, *x*_*v*_) under the hypotheses of uncorrelated sampling and independence: *p* (*x*_*u*_, *x*_*v*_) = *p* (*x*_*u*_) *p* (*x*_*v*_) . Hence, we can be more confident that pairs of conformations which we do not observe simultaneously have some energetic conflict one another (rather than unobserved by mere chance) than we would be with no coarse-graining; controlling chance fluctuations in low-probability states tames a major source of error in entropy estimation (*96*).

The output of this step is a set of conformations, C, which is a partition of the von Mises mixture components. Each observation, θ (*t*), is classified to the conformation with the maximum likelihood of generating it, which is the sum of the likelihoods for all mixture components mapped to that conformation.

#### Maximum likelihood tree inference

If we have a distribution, *p* (*X*_1_…*X*_*L*_), which is not necessarily a tree, then the tree distribution *p*_*T*_ minimizing the information loss (called the Kullback-Leibler divergence, *D*_*KL*_) from this distribution is identified with the spanning tree of *X*_1_…*X*_*L*_ with the maximum mutual information, which can be shown with some algebra. Writing *S* (*X*_1_…*X*_*L*_) for the joint entropy of the residues, *S* (*X*_*u*_) for the marginal entropy of residue *u*, and *I* (*X*_*u*_, *X*_*v*_) for the mutual information between residues *u* and *v*, one has:

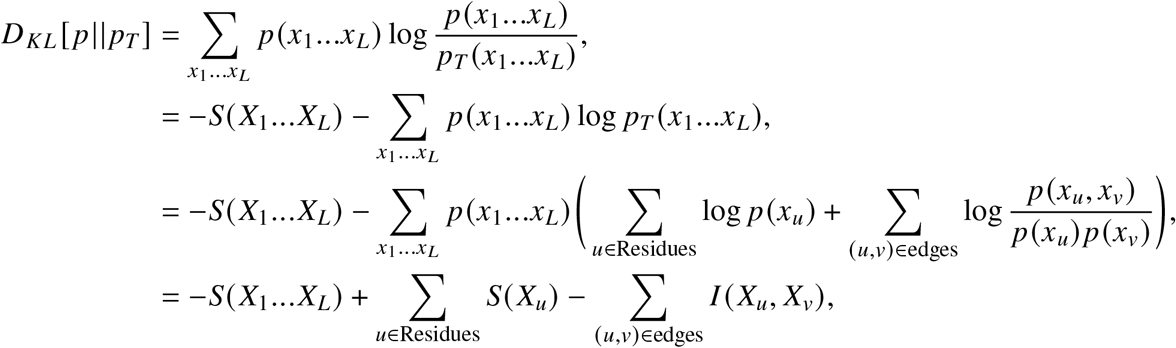

where the last line follows from the fact that the single and pairwise marginals of *p*_*T*_ are equal to those of *p*. The first two terms on the last line are the sum of the marginal entropies, *S* (*X*_*u*_), of the residues minus the true entropy of *p*. The only term that depends on the structure of the tree is the last one which is the summed mutual informations. The tree which minimizes the information loss from the true distribution is therefore the one which maximizes the sum of mutual informations. Moreover, the Kullback-Leibler divergence is the expected difference in log-likelihood per observation between the true distribution and the tree; therefore, the tree that generates the joint distribution closest to the true distribution is the maximum information spanning tree whose component distributions, { *p* (*x*_*u*_)} and { *p* (*x*_*u*_, *x*_*v*_)}, are equal to their counterparts in the distribution *p* (*x*_1_…*x*_*L*_).

#### Posterior distribution of the entropy

Optimizing the likelihood of a mixture model using the expectation-maximization algorithm does not lead to a global optimum, but rather to a fixed point of the update rule determined by the randomly initialized parameters. Random variation in the mixture model parameters propagates through all subsequent steps in the analysis to produce random variation in the final estimate of the conformational entropy. We therefore derived a heuristic method for estimating the distribution of plausible conformational entropies from a single tree which encapsulates this uncertainty. Although our treatment of uncertainty is somewhat heuristic, we find empirically that it captures the variability associated with random initializations.

We model the posterior distribution of the entropy conditional on a single tree, T, using a normal distribution,

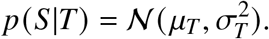

##### Posterior mean

The posterior mean, *µ*_*T*_, is the expected entropy of the tree distribution given the observed counts {**n**_*u*_}, {**n**_*uv*_} of single-residue and residue-pair states for branches included in the tree. This is just the sum of expectations of residue entropies and branch mutual informations. We compute these expectations with respect to Dirichlet distributions parameterized by the counts. The expected value of the entropy of a realization of a Dirichlet distribution parameterized by a vector of counts, **n**, is given by the expression (*97*):

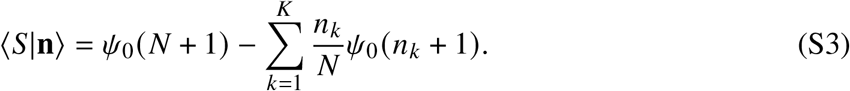

where *n*_*k*_ denotes the *k*-th state, *K* is its dimension, *N* is the number of states, *N* is the total number of observations, and *ψ*_0_(∗) is the polygamma function of order zero (also called the digamma function), defined as 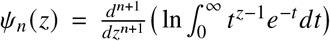. It is straightforward to then compute the expected mutual informations as

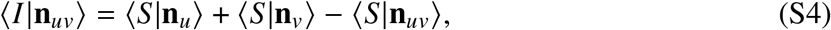

leading to the expression for the posterior mean entropy of the tree:

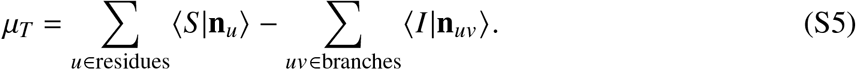

##### Posterior variance

In estimating the uncertainty of the entropy we hope to capture the expected squared deviation of *µ*_*T*_ from its (unknown) true value, which is the value that would be obtained by initializing the program many times and averaging. Our posterior variance aims to account for three sources of uncertainty 1) fluctuations in the number of observed counts for single and pairwise residue states that arise due to finite sampling 2) uncertainty due to the use of the use of the maximum likelihood estimator in determining the tree structure 3) uncertainty about the entropy that arises due to uncertainty about the optimal tree structure itself. This led us to the following decomposition of the variance:

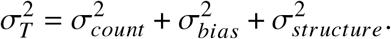

The counting variance is accounted for using the following expression for the expected squared entropy of a realization from a Dirichlet distribution (*97*),

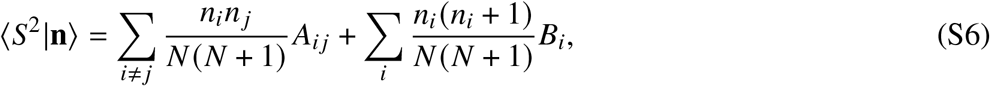

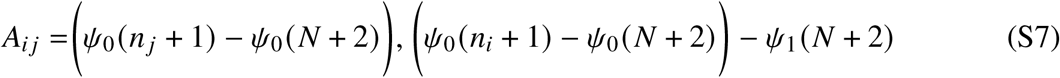

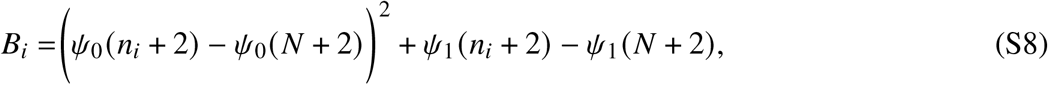

from which the variance is obtained as ⟨*S*^2^|**n**⟩ − ⟨*S*|**n**⟩ ^2^ . We approximate the counting fluctuations in the residue entropies and mutual informations as independent, so that their contributions to the total variance are simply additive.

We estimate the bias component of the expected squared deviation as

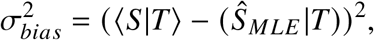

where 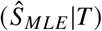 is the entropy of the tree distribution when each residue entropy and mutual information are calculated using the plug-in estimator 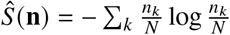.

We emphasize that we do not attempt to make explicit bias corrections, but only to obtain an estimate of its contribution to the mean-squared error. We give a few justifications for this. First, we observed that the estimated biases for the individual entropies and mutual informations are typically not much larger than the estimated standard deviations, which led us to expect that approximating the bias as a component of the variance might lead to more accurate interval estimates than attempting an explicit correction. Second, the true shape of the posterior distribution of the entropy is unknown, so incorporating an estimate of bias into the uncertainty reflects some degree of uncertainty about which estimator is most appropriate given the data without imposing any prior (or hyperprior) distribution on the counts vector. Third, and most importantly, our broader goal is to accurately capture differences in entropy, as total protein entropies are not observable. Towards this end, we considered explicit bias correction to be undesirable, as it is not generally possible to distinguish between bias that due to differing numbers of states and a genuine entropy difference due to a change in the number of occupied free energy wells. We therefore judged it more appropriate to account for bias as a source of uncertainty than to attempt corrections.

The structural uncertainty aims to capture the variation in the entropy that originates from changes in the structure of the maximum likelihood tree that arise from different partitions of the configurational landscapes of residues, which result in correlated differences in the presence of branches connecting distinct residues. In doing this, we also account, to some extent, for the possibility of spurious mutual information between residues that arises due to undersampling of slow degrees of freedom. Let 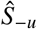 denote the entropy estimate obtained by deleting the *u*-th residue and estimating the entropy by applying the Chow-Liu algorithm (*63*) to the remaining residues and 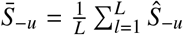 denote the empirical average of these replicates. We estimate the structural variance as the empirical variance of all replicates formed this way:

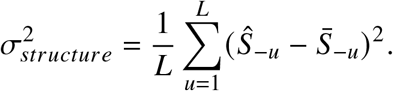

In spite of the heuristic nature of this decomposition, the credibility intervals produced this way appear to have good frequentist coverage in our sample, with 93.7% of the single-tree 95% credible intervals containing the average entropy over 21 fits for a representative set of eight thermodynamic states (Supplemental figure S7).

#### Bayesian model for allosteric mutation predictions

We extracted the number of allosteric mutations at each site from the supplemental data provided in (*58*). We adopted the definition of allosteric mutation used by the authors: a mutation whose effect on CRIPT-binding was significantly greater than the effect of a random binding site mutation on CRIPT-binding. We modeled the number of allosteric mutations at each site using a binomial distribution,

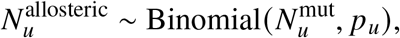

where 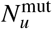 is the total number of point mutations reported at site *u* in the sequencing data (*58*), 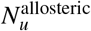 is the number of allosteric mutations reported at this site, and *p*_*u*_ is a latent variable which is our estimated probability that a random mutation at site *u* will be allosteric.

For most residues, all 19 possible mutations were observed, but for a minority of residues only 17 or 18 mutant alleles were observed. This heterogeneity in allele counts is a significant source of uncertainty, as we cannot know for certain the reason that alleles are absent from the sequencing data. This also necessitates a hierarchical model because the probability of an allosteric mutation cannot be estimated in terms of a binomial regression with one fixed number of observations. To account for this, we modeled *p*_*u*_, the site-specific probability of allosteric mutation as a random draw from a beta distribution. The site-specific predictors are given by

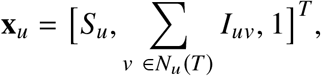

where *S*_*u*_ is the marginal entropy of residue 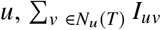 is the sum of all mutual informations between residue *u* and its neighbors *N*_*u*_ *T* in the tree, and the final term represents an intercept. These features are related to our estimate of the number of allosteric mutations according to

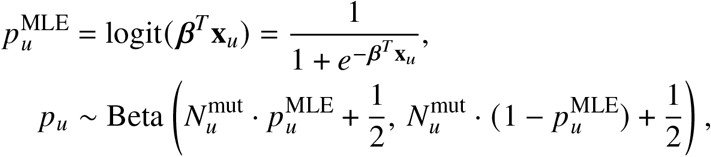

where *β* is the vector of model coefficients. This is similar to a logit-binomial regression model, which would be obtained by setting 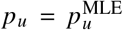 and 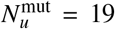. Instead, we have replaced the maximum likelihood step by simulating a Bayesian inference step using a Beta (Jeffreys) 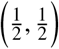 prior. The use of the Beta distribution allows us to integrate over uncertainty in *p*_*u*_ that arises due to unobserved alleles while ensuring that uncertainty increases with the number of unobserved alleles.

We used a multivariate normal distribution as a prior for our coefficient vector *β* with a diagonal covariance matrix and a weakly-informative Half-Cauchy prior with scale parameter equal to 25 for the variance. The model was estimated using Markov Chain Monte Carlo (MCMC) with 5000 draws using the No U-turn sampler (*98*).

#### ciMIST software dependencies

ciMIST uses MDTraj (*99*) to process trajectories and nerfax (*100*) to handle conversion from internal to cartesian coordinates. JAX (*101*) is used for numerical calculations and NetworkX (*102*) for graph algorithms.

#### Experimental data sourcing

ITC data was obtained from the original publications (*28, 73*), with experimental uncertainties for the ER*α* data obtained from the authors. NMR and HDX data were scraped from graphs in the original publications (*12, 13, 28*) using the WebPlotDigitizer tool. Experimental structures used to create starting structures for MD simulations were obtained from the PDB.

#### Data Visualization

Plots were made in Python using Matplotlib (*103*). Protein visualization was done in PyMOL (*104*). Colormaps displayed on proteins are drawn from the cmocean libary (*105*).

## Supplementary Text

**figure S1:**
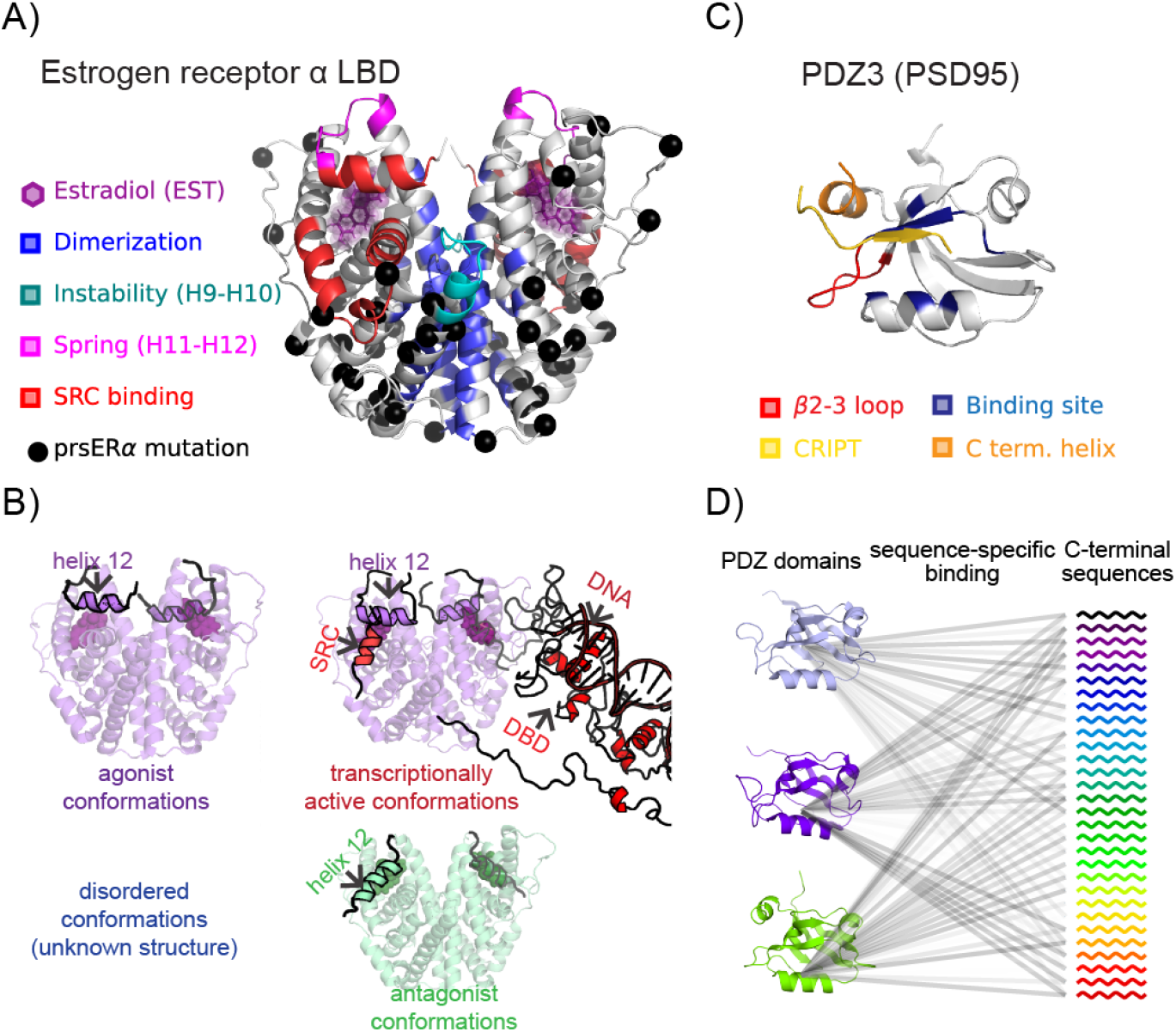
Allostery in estrogen receptor *α* and PDZ3. A) Estrogen receptor *α* ligand-binding domain (LBD, PDB: 1QKU) homodimer bound to estradiol. Sites colored in blue, teal, pink, and red influence activities from outside the hormone binding site. Sites shown in black spheres are those at which stabilizing mutations were introduced to create an engineered thermostable estrogen receptor. B) Major conformations of ER*α*. Binding of estradiol (purple) in the ligand-binding pocket stabilizes homodimers with an inward-facing H12 conformation poised for binding to steroid receptor coactivators (SRCs), leading to the formation of an active transcriptional complex on DNA estrogen response elements. Binding of selective estrogen receptor modulators (SERMs; green) induces an H12 conformation oriented away from the homodimer interface, precluding the binding of SRCs. The unliganded state is thought to be disordered, but is not known. Structures displayed are based on PDB entries 1QKU (agonist), 3ERT (antagonist), and SASBDB entry SASDDU8 (DNA bound). C) Structure of the third PDZ domain (PDZ3, PDB: 5HEB) from PSD95. The CRIPT peptide, binding site, and allosteric sites are labelled. D) Schematic illustrating an interaction network showing distinct profiles of specificities of PDZ domains for C-terminal peptide sequences Models shown are based on PDB entries 1KEF, 3ZRT, and 5HEB (top to bottom).

**figure S2:**
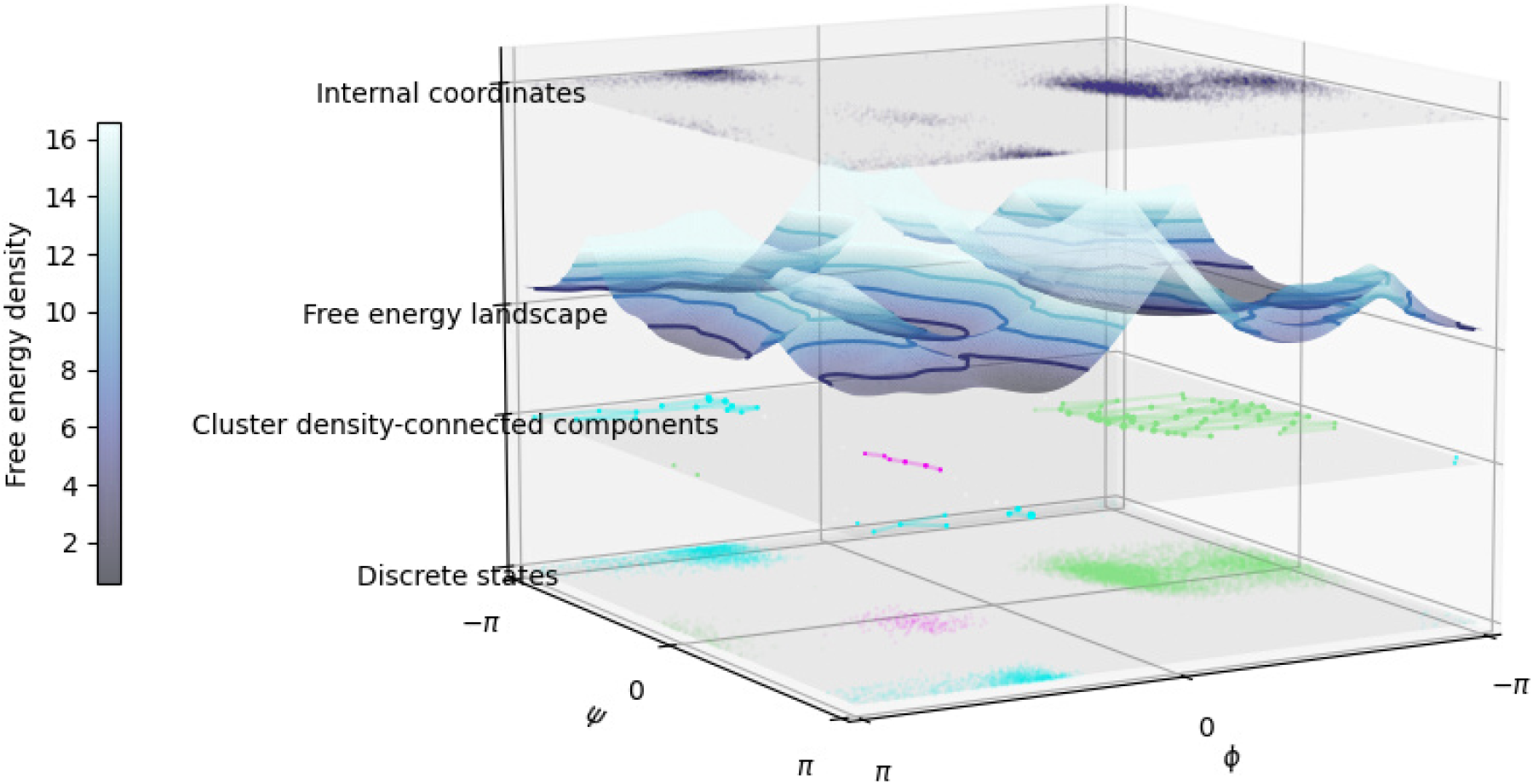
Example of the application of our conformational inference algorithm projected onto the *ϕ* and *ψ* angles of a glycine. The top row shows a scatter plot of the observed angles. The second row shows the free energy landscape estimated by the von Mises mixture model. The third shows a clustering of the 100 von Mises mixture components by a density-overlap based distance. The bottom row shows the final labeling of all observations in the trajectory.

**figure S3:**
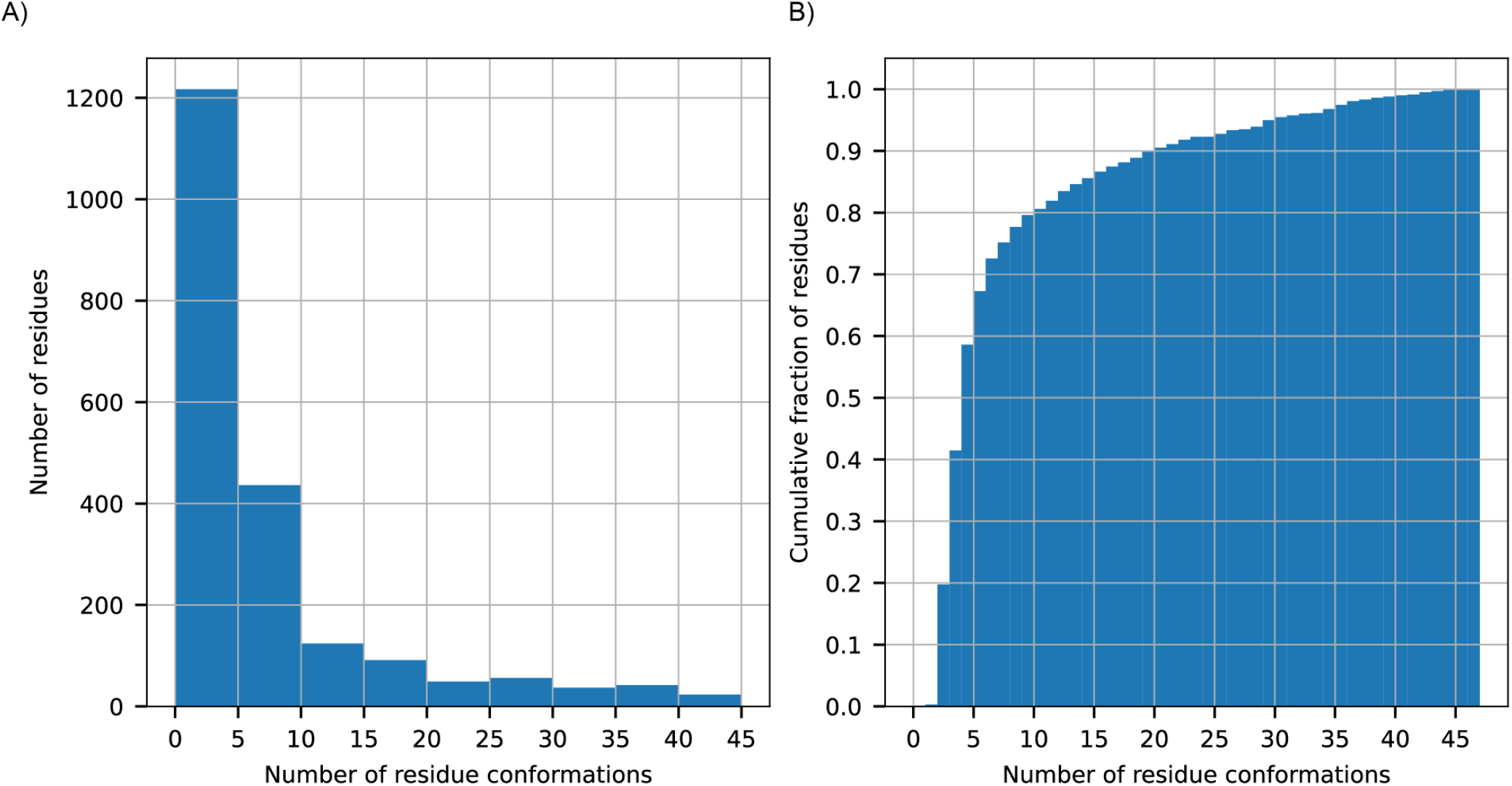
Frequencies of numbers of residue conformations across all ensembles analyzed in this work. A) Histogram showing the absolute number of times our conformational inference algorithm inferred different numbers of residue conformations across all ensembles analyzed in this work. Note we use a bin width of five for the histogram. B) Cumulative fraction of times our conformational inference algorithm inferred a number of residue conformations less than or equal to a given value. Note the bin width (x-axis) is one.

**figure S4:**
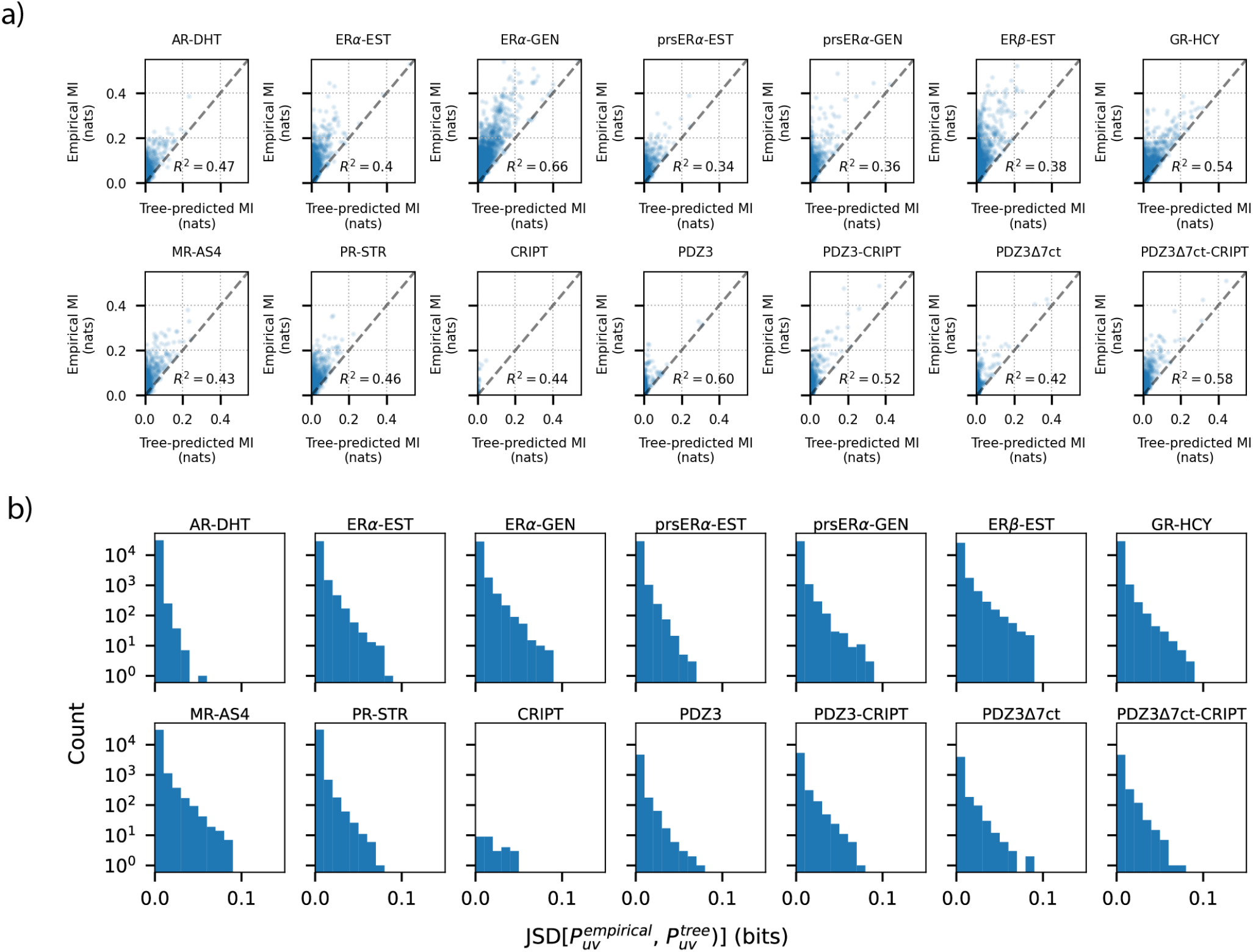
Consistency of tree-predicted pairwise conformational distributions with empirical conformational distributions for pairs of residues not retained in the tree. A) Correlation between tree-predicted and empirical (maximum likelihood) mutual information for edges not included in the maximum information spanning trees for all systems analyzed in this work. B) Histograms showing frequecies of Jensen-Shannon divergences (JSD) between tree-predicted and empirical pairwise conformational distributions. JSDs are expressed in units of bits so that the maximum possible value is 1.

**figure S5:**
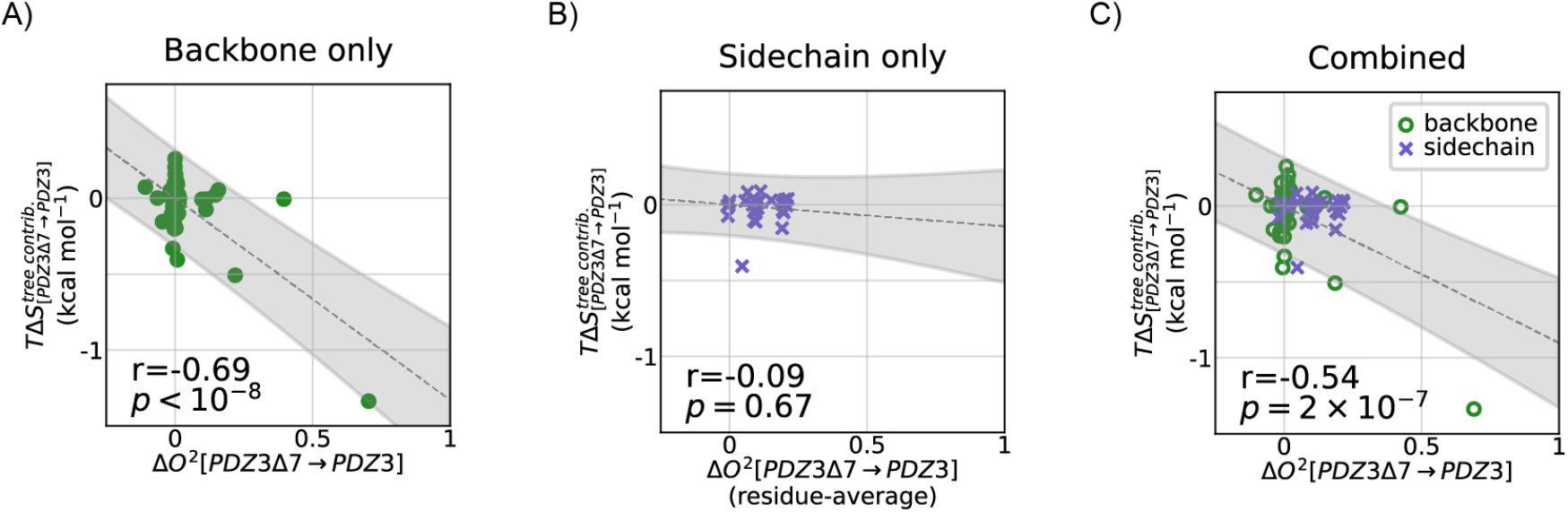
Correlation of residue entropy contributions with NMR. A) Correlation between differences in residue entropy contributions and changes in backbone *O*^2^order parameters (*28*) between PDZ3 and PDZ3Δ7-ct. B) Correlation between differences and residue entropy contributions and changes in side-chain *O*^2^order parameters (*28*) between PDZ3 and PDZ3Δ7-ct. C) Correlation between differences and residue entropy contributions and the combined set of *O*^2^order parameters (*28*) between PDZ3 and PDZ3Δ7-ct. Side-chain order parameters are residue-averaged.

**figure S6:**
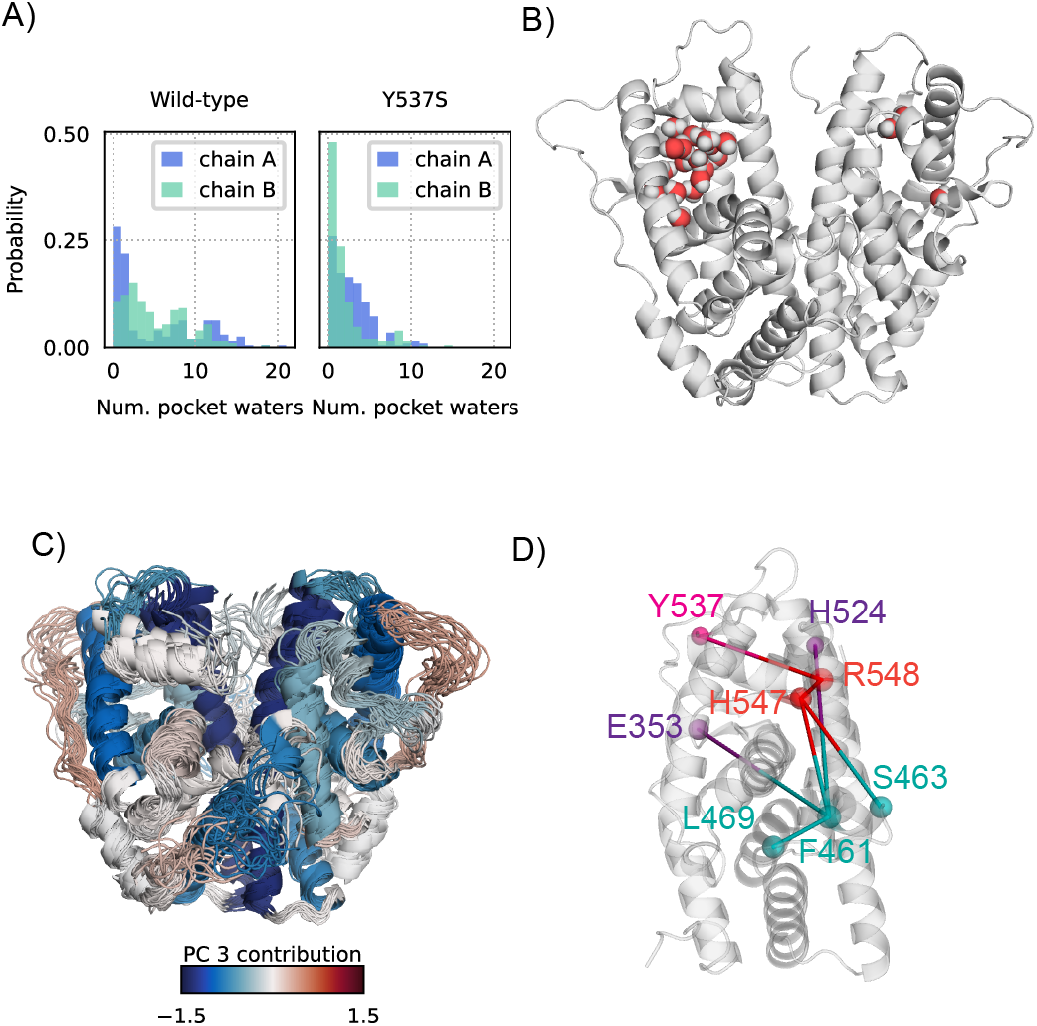
A) Histograms of the number of waters in the binding pocket as defined by the convex hull of its atoms in each frame. Differences between the wild-type and Y537S pocket hydrations are significant with *p* < 10^−6^ . B) A snapshot of the wild-type apo state showing a lifted H12 conformation and hydrated binding pocket. C) 20 random snapshots drawn from our sampled conformational ensemble for the wild-type apo state, with secondary structural elements colored by their contribution to the third principal component, which represents the ordering of the receptor common to all ligand binding. Blue regions are generally ordered by ligand-binding, while red ones are disordered. D) Paths in the ciMIST tree connected the H9-H10 region, H12, Y537, and two binding pocket residues which form hydrogen bonds with estradiol.

**figure S7:**
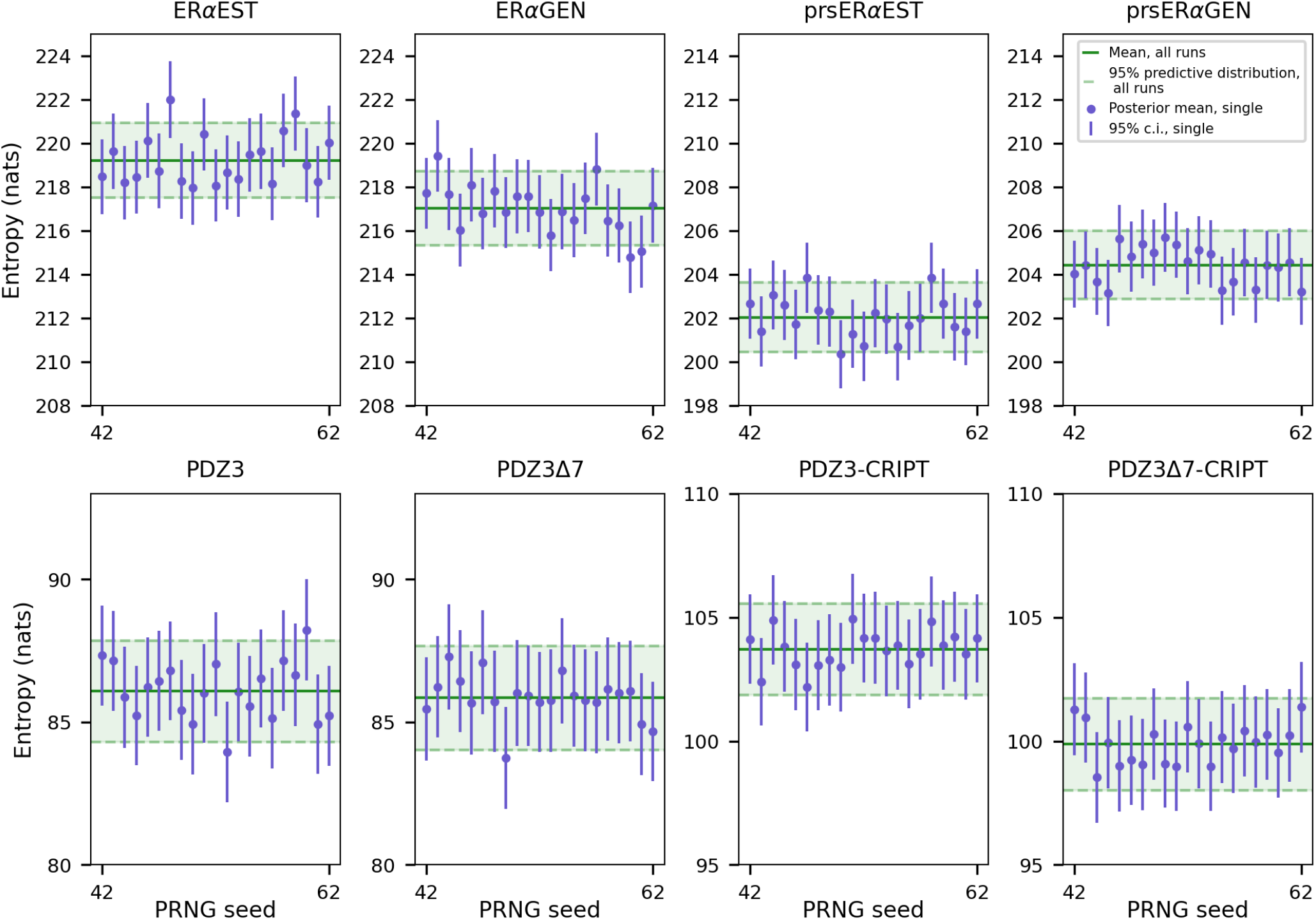
Uncertainty quantification for tree entropies. Shown are the average entropies obtained from running the program with 21 different pseudo-random number generator (PRNG) seeds, the 95% posterior predictive distribution estimated from these runs and the posterior means and credibility intervals from the each run.

**figure S8:**
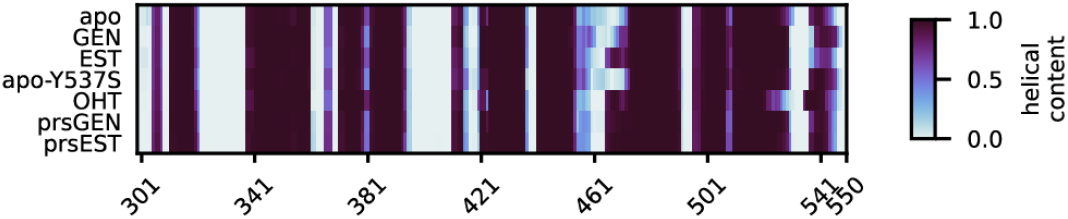
Helical content as quantified using the DSSP algorithm in simulations for each ER*α*-LBD state.

**figure S9:**
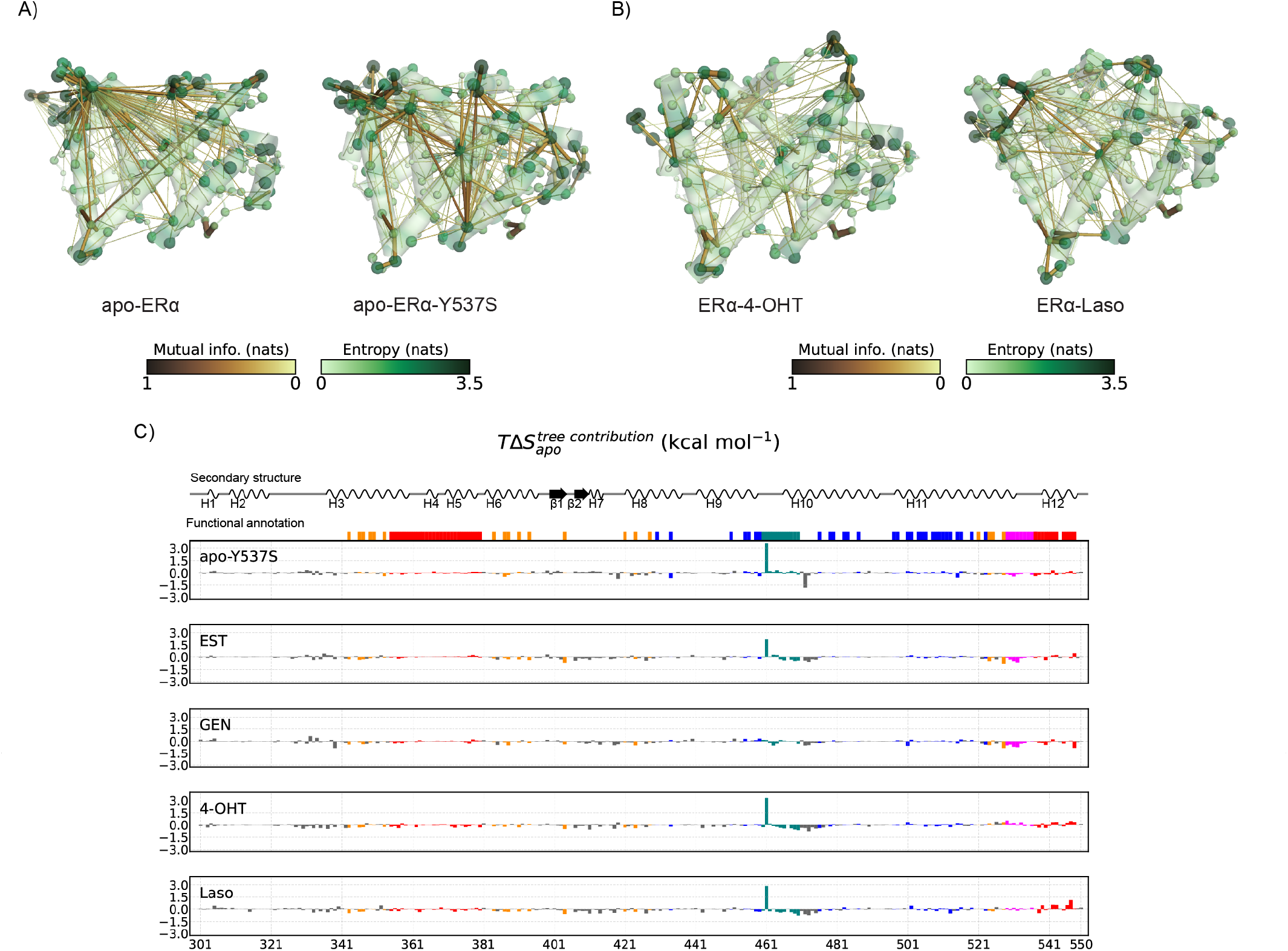
a) Maximum information spanning trees for apo states of wild-type ER*α* and the Y537S variant. b) Maximum information spanning trees for ER*α* bount to anti-estrogen ligands 4-hydroxy-tamoxifen (4-OHT) and lasofoxifene (Laso). c) Differences in residue entropy contributions from the wild-type apo state of ER alpha for the Y537S mutant and all liganded states.

**Table S1:** Details of molecular dynamics simulations. State refers to the thermodynamic state that we modelled. PDB refers to the PDB code of the model(s) used to create the starting structure. Parentheses demarcate structures used as templates for a homology models in Modeller (m) or ColabFold (a) (*106*). Other than for the Y537S and prsER*α* simulations, mutations were made to restore the UniProt sequence corresponding to the PDB id. Residues not present in PDB structures shown were modelled using “molecular dynamics”-based refinement using the Modeller software.

| State | PDB | Residues | Total time (μs) | Replicates |
| --- | --- | --- | --- | --- |
| PDZ3 | 1BFE | 302-402 | 10.5 | 6 |
| PDZ3Δ7ct | 1BFE | 302-395 | 10.5 | 6 |
| PDZ3Δ7ct-CRIPT | 5HEB | 302-395 (A), 1-9 (B) | 10.5 | 6 |
| PDZ3-CRIPT | 5HEB | 302-402 (A), 1-9 (B) | 10.5 | 6 |
| CRIPT | 5HEB | 1-9 (B) | 10.5 | 4 |
| ERα (apo) | (2B23,4Q13)<sup>m</sup> | 301-550 (A, B) | 5.1 | 6 |
| ERα-C3D | 6VJD | 301-550 (A, B) | 4.9 | 6 |
| ERα-EST | 1QKU | 301-550 (A, B) | 5.5 | 6 |
| ERα-GEN | 2QA8 | 301-550 (A, B) | 5.3 | 6 |
| ERα-OHT | (3ERT)<sup>a</sup> | 301-550 (A, B) | 5.3 | 6 |
| ERα-Y537S (apo) | 2B23 | 301-550 (A, B) | 5.1 | 6 |
| prsERα-EST | 7NEL | 302-548 (A, B) | 5.3 | 6 |
| prsERα-GEN | 7NFB | 302-548 (A, B) | 5.7 | 6 |
| AR-DHT | 1T7T | 669-920 | 5.3 | 4 |
| ERβ-EST | 3OLS | 261-500 (A, B) | 4.0 | 4 |
| GR-HCY | (4P6X)<sup>m</sup> | 529-776 | 6.1 | 4 |
| MR-AS4 | 2AA2 | 737-984 | 6.5 | 4 |
| PR-STR | 1A28 | 678-933 | 5.5 | 4 |

